# A neuroskeletal atlas of the mouse limb

**DOI:** 10.1101/2020.09.18.303958

**Authors:** Madelyn R. Lorenz, Jennifer M. Brazill, Alec Beeve, Ivana Shen, Erica L. Scheller

## Abstract

Nerves in bone play well-established roles in pain and vasoregulation and have been associated with progression of skeletal disorders including osteoporosis, fracture, arthritis and tumor metastasis. However, isolation of the region-specific mechanisms underlying these relationships is limited by our lack of comprehensive maps of skeletal innervation. To overcome this, we mapped sympathetic adrenergic and sensory peptidergic axons within the limb in two strains of mice (B6 and C3H). In the periosteum, these maps were related to the surrounding musculature, including entheses and myotendinous attachments to bone. Locally, three distinct patterns of innervation (Type I, II, III) were defined within established sites that are important for bone pain, bone repair, and skeletal homeostasis. In addition, we mapped the major nerve branches and areas of specialized mechanoreceptors. This work is intended to serve as a guide during the design, implementation, and interpretation of future neuroskeletal studies and was compiled as a resource for the field as part of the NIH SPARC consortium.

---

Peripheral nerves target nearly every organ in the body to communicate afferent and efferent signals to and from the brain; the skeleton is no exception^1^. Neurons in and around the bone have well-established roles in the generation and maintenance of pain associated with skeletal pathologies including fracture, tumor progression, and arthritis^2–4^. Within cortical bone and bone marrow, sensory and sympathetic nerve fibers track with the arteriolar vasculature^5–7^ and coordinate vasoregulatory responses^8–11^. Additional functions have also been proposed, including modulation of bone formation and turnover in the context of skeletal development^12^, adaptation to mechanical loading^13,14^, repair, and regeneration^15–17^, as well as regulation of bone marrow adipogenesis^18^, joint disease^19^, and control of hematopoietic cell maturation and egress^20^.

Direct neural modulation of skeletal function is dependent upon the precise localization of axons near target cells and the capacity of the neurons to sense the bone microenvironment and/or to release neurotransmitters. Sensory peptidergic axons, often marked by calcitonin gene-related peptide (CGRP), as well as sympathetic axons containing tyrosine hydroxylase (TH), the rate-limiting enzyme in catecholamine synthesis, are distributed in and on the bone^6,21^. These neuronal subtypes have been shown to innervate the periosteum in a meshwork pattern, traverse the cortical bone through nutrient canals, and to innervate the marrow cavity^6^. Skeletal axon varicosities, presumed sites of neurotransmitter release and receptor clustering, have been observed in close proximity to bony surfaces and resident osteoblasts, osteoclasts, and their precursors, as well as at sites of marrow adipose tissue and hematopoietic cells^21,22^. While the morphology and molecular phenotype of skeletal axons have been well studied^23^, the inherent challenges of visualizing and quantifying neural fibers in calcified tissue have led to a functional and anatomical neuroskeletal map that is incomplete. This is complicated by the heterogeneity of the bone architecture, the complex tissue interactions therein, and the opposing influences of the sensory and sympathetic arms of the nervous system that converge to influence the physiological functions of the skeleton^1^.

To begin to overcome this, we used thick section immunohistochemistry and tissue clearing, in concert with high-resolution imaging and axon quantification techniques, to map skeletal innervation in the lower limb with emphasis on comprehensive mapping of axons within and around the femur and tibia. In addition, we defined the location, prevalence, and morphology of specialized mechanoreceptor endings. This information has been compiled into a reference atlas of skeletal innervation that can be used to inform future neuroskeletal studies. This resource was generated as part of the SPARC consortium (Stimulating Peripheral Activity to Relieve Conditions), a call by the National Institutes of Health in the United States to generate foundational neuroanatomic maps of axons within peripheral tissues with the aim of supporting the development of novel neuroceuticals for targeted treatment of pain and other conditions.

## RESULTS

### Validation of neuroskeletal immunoassay and axon quantification in the C3H and B6 mouse tibia

Prior to creating the atlas, we validated our axon tracing and immunostaining paradigm relative to previous publications (Table 1)^6,24,25^. To achieve this, we quantified CGRP+ sensory and TH+ sympathetic axons in the periosteum and bone marrow of 12-week-old male C57BL/6J (B6) and C3H/HeJ mice (C3H) at four levels along the length of the tibia. Region-specific bone parameters including cortical and trabecular bone volume fraction (BVF) were also assessed. Levels of the tibia selected for analysis were distinguished by bone morphology at the metaphysis, mid-diaphysis with tibial ridge, the diaphysis proximal to the tibia-fibula junction (TFJ), and the diaphysis just distal to the TFJ. Analyzed levels approximated to 10% (L1), 30% (L2), 50% (L3), and 60% (L4) distal to the knee for B6 mice (Fig 1.a,b). Consistent with previous reports^26,27^, the diaphyseal cortical BVF was significantly higher in C3H than B6 mice (L2, L3; Fig.1c). However, cortical BVF was matched in the two strains at both the proximal and distal metaphysis (L1, L4; Fig.1c). Trabecular BVF was comparable in both strains at the proximal metaphysis (L1; Fig.1g), also consistent with prior comparisons at this region and age^28^. Trabecular bone was largely absent in the tibial diaphysis of both B6 and C3H mice (L2-4).

**Figure 1.**
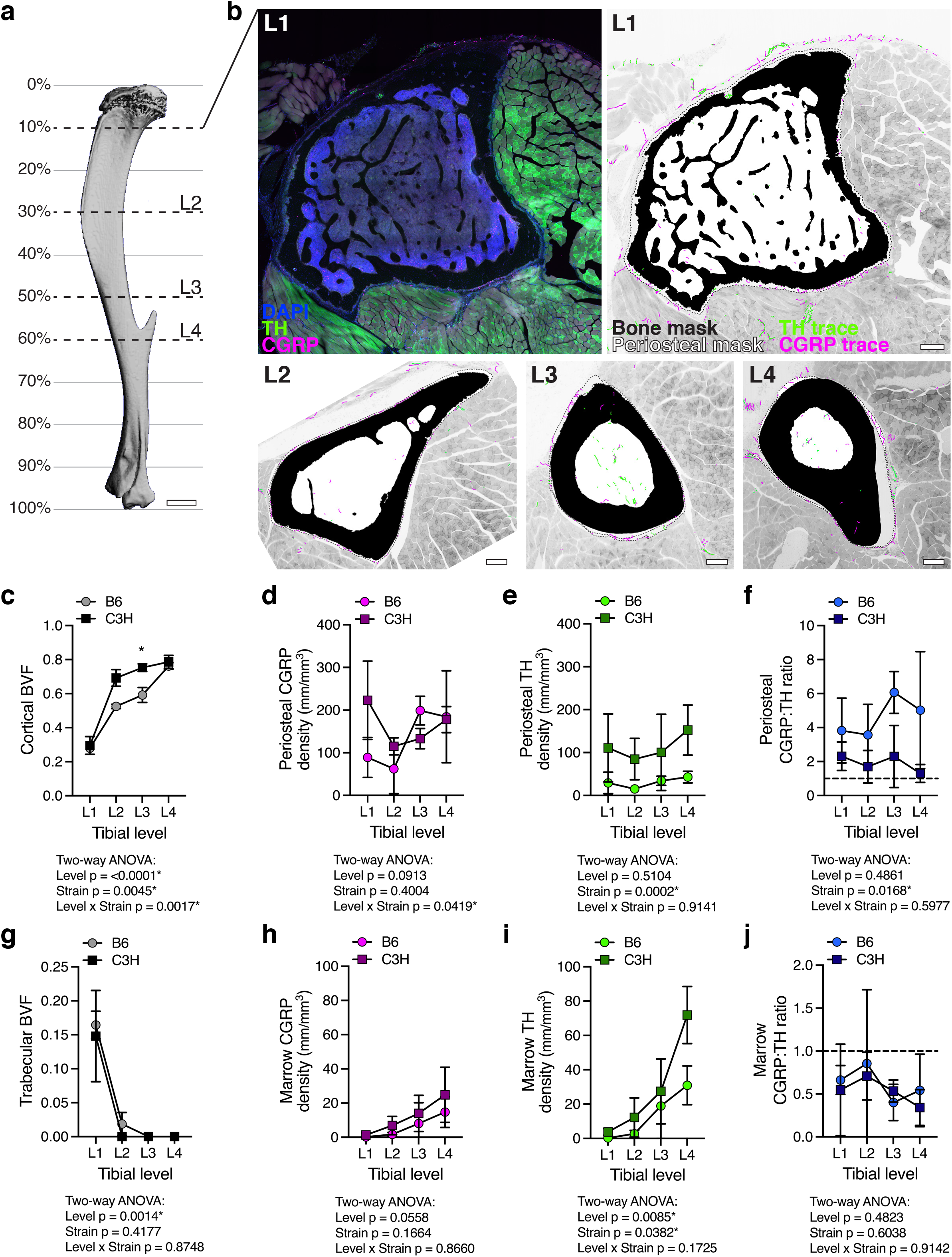
Neuroskeletal axon density varies by bone compartment, tibial level, and mouse strain. (**a**) 3D rendering of a μCT scan of a 12-week-old male B6 tibia, lateral view. L1-4 represent levels analyzed as distinguished by cross-sectional bone morphology. Percentages demarcate the approximate distance from the knee as a percentage of total tibia length. Scale bar, 1 mm. (**b**) L1 Left: Representative confocal micrographs through a 50-μm-thick transverse cross section with immunolabeled CGRP+ sensory axons (magenta) and TH+ sympathetic axons (green), as well as DAPI+ nuclei (blue). L1 Right and L2-L4: Bone mask (black), periosteal mask (dotted line), CGRP+ axon traces (magenta) and TH+ axon traces (green) overlaid on a grayscale max projection to visualize tibial innervation in the context of the musculoskeletal system at the metaphysis (L1), the diaphysis with tibial ridge (L2), the diaphysis proximal to the TFJ (L3), and distal to the TFJ (L4). Scale bars, 200 μm. (**c-j**). Quantification of compartmentalized bone volume fraction (BVF) and innervation density from B6 and C3H mice. (**c**) BVF of cortical bone. (**d,e**) CGRP+ and TH+ axon density (mm/mm^3^) within the overlying periosteum. (**f**) Periosteal CGRP:TH ratio. (**g**) BVF of trabecular bone. (**h,i**) CGRP+ and TH+ axon density (mm/mm^3^) within the marrow cavity. (**j**) Marrow CGRP:TH ratio. Data represent the mean ± SD; n = 3 for each strain and level; Two-way ANOVA with Sidak’s multiple comparisons test; * p < 0.050

**Table 1.** Previous quantification of TH+ and CGRP+ axon density in mouse bone.

| Publication | Mouse Information | Bone | Compartment | Immunostain | Summary of validated results |
| --- | --- | --- | --- | --- | --- |
| Mach et al., 2002 | C3H/HeJ male<br>6-7 weeks, 20-25 g | Femur<br>proximal head,<br>diaphysis,<br>distal head | Periosteum,<br>cortical bone,<br>bone marrow | CGRP<br>TH | • Nerve density: periosteum > bone marrow<br>• In periosteum, CGRP>TH (1.3 to 2.4-fold)<br>• In bone marrow, CGRP<TH (0.3 to 0.5-fold) |
| Castañeda-Corral et al., 2011 | C3H/HeJ male<br>20-25 g | Distal femur | Periosteum,<br>cortical bone,<br>bone marrow | CGRP | • Nerve density: periosteum > bone marrow<br>• TH not assessed |
| Chartier et al., 2018 | C57Bl/6J male<br>10 days, 3 months,<br>24 months | Distal femur | Periosteum,<br>bone marrow,<br>cortical pores | CGRP<br>TH | • Nerve density: periosteum > bone marrow<br>• 3-month periosteum CGRP>TH (1.3-fold)<br>• 3-month bone marrow CGRP<TH (0.7-fold) |

At each level, sensory and sympathetic axons were traced and the total fiber length within each compartment was expressed relative to the total volume (Fig.1b). As reported previously for femur (Table 1), the periosteum was the most densely innervated compartment of the tibia for both B6 and C3H mice. Within the periosteum, we found that CGRP+ sensory axons demonstrated level-specific densities that were dependent on mouse strain (Fig.1d). By contrast, TH+ sympathetic axon density was relatively constant along the length of the periosteum. C3H mice had a greater absolute TH+ fiber density at all levels relative to B6 (Fig.1e). Independent of strain, periosteal CGRP+ sensory fibers were more prevalent than TH+ sympathetics at all levels of the tibia (periosteal CGRP:TH ratio > 1; Fig.1f). This ratio was higher in B6 animals than C3H (Fig.1f). Within the marrow cavity, innervation density increased from proximal to distal independent of strain (Fig.1h,i). Converse to the ratio of neural subtypes observed in the periosteum, TH+ sympathetic innervation consistently exceeded CGRP+ sensory innervation in all regions of the bone marrow (CGRP:TH ratio < 1; Fig.1j). Overall, our immunolabeling and analysis paradigm corroborates previous reports of compartmentalized innervation density (Table 1), and additionally demonstrates that this varies by strain. Our results further establish a gradient of bone marrow axon density that increases from proximal to distal along the length of the tibia.

### Identification of three distinct patterns of periosteal innervation (Type I, II, III)

Next, we applied this validated immunolabeling and tracing paradigm to assess the periosteal axon profiles throughout the entire length and circumference of the tibia and femur of male and female 12-week-old C3H and B6 mice. The periosteum is the layer of connective tissue that envelops the bone and provides a supportive microenvironment for vasculature, nerves, and periosteal cells^29,30^. We identified three consistent patterns of adult periosteal innervation that aligned with the attachments formed between the periosteum and the surrounding tissues, independent of mouse sex or strain. Within each innervation pattern, we observed variations in local axon density and orientation that should be considered for future studies pertaining to the relationships between nerves and bone, as described below.

#### Type I: Aneural

Many regions of the bone surface were aneural, completely lacking CGRP+ sensory or TH+ sympathetic innervation (Fig.2a-d). Aneural regions were observed primarily where tendons or ligaments interfaced with the bone at sites called entheses. There are two main classes of entheses: fibrocartilaginous and fibrous^31,32^. Fibrocartilaginous entheses comprise continuous zones connecting tendinous fibers through unmineralized and mineralized fibrocartilage tissue that blend imperceptibly into bone (Fig.2a). At fibrous entheses, the tendon is continuous with a thick fibrous layer of the periosteum (Fig.2b). The only axons found in these regions were located in the fascial tissue lining the tendon or ligament, often referred to as the epitenon. Type I regions greatly contributed to the discontinuous nature of periosteal innervation.

**Figure 2.**
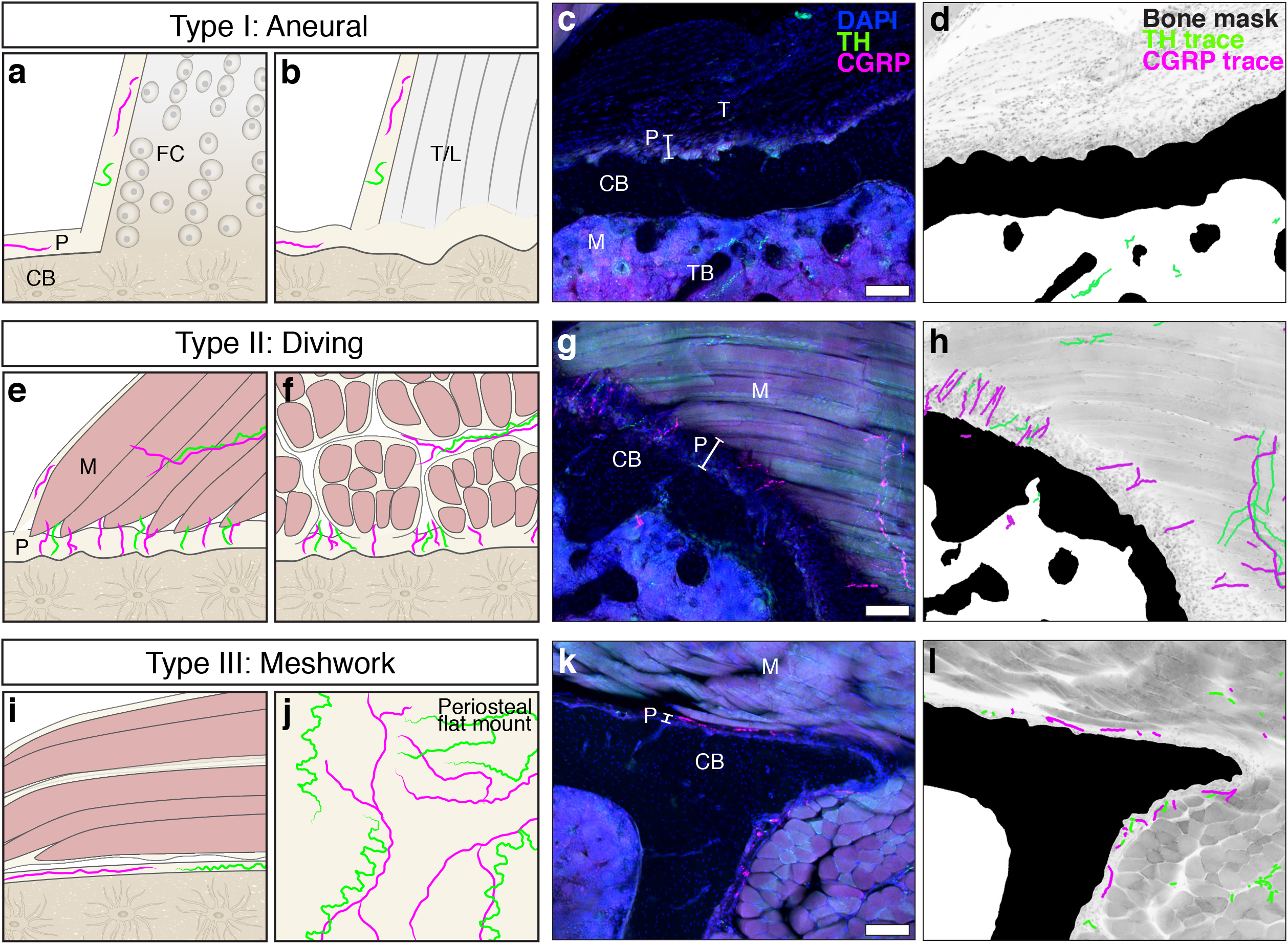
Peripheral axons innervate the periosteum in three distinct patterns (Type I, II, III). (**a-d**) *Type I: Aneural* patterns are found at entheses, sites where tendons or ligaments attach to the cortical bone (CB) surface. Aneural regions of the periosteum are devoid of CGRP+ (magenta) and TH+ (green) axons. (**a**) Tendinous fibers connect seamlessly to bone through unmineralized and mineralized fibrocartilage tissue (FC) at fibrocartilaginous entheses. (**b**) The tendon or ligament (T/L) is continuous with a thick fibrous layer of the periosteum (P) at fibrous entheses. (**c**) Confocal micrograph through a 50-μm-thick transverse cross section a tibia from a 12-week-old C3H male with immunolabeled CGRP+ sensory axons (magenta) and TH+ sympathetic axons (green), as well as DAPI+ nuclei. (**d**) Corresponding neuroskeletal traced/masked analysis overlaid on a grayscale max projection demonstrating an aneural enthesis at the tibia metaphysis where the semimembranosus tendon attaches to the bone. Trabecular bone (TB) is visible in the marrow cavity (M). (**e-h**) *Type II: Diving* innervation patterns are found at sites where axons transit directly from the associated muscle tissue through the fibrous and cambium layers of the periosteum at a 90-degree angle to the cortical bone surface as visualized here in the C3H tibia where the popliteus muscle attaches to the bone. (**i-l**) *Type III: Meshwork* innervation patterns are found at sites where fascia attach to bone. Axons are woven through the thin periosteal layer at these sites and run parallel to the bone surface as evident in cross-section (**i,k,l**). The mesh-like network of periosteal axons at these sites is clearly visualized by periosteal whole-mount preparation (**j**, see ^7,34^). Scale bars, 100 μm.

#### Type II: Diving

In other regions, axons transited directly from the associated muscle tissue through the fibrous and cambium layers of the periosteum at a 90-degree angle to the cortical bone surface (Fig.2e-h). Type II regions contained TH+ sympathetic axons and an abundance of CGRP+ sensory axons that appeared to dive from the sarcolemma or muscle fascia to the bone surface. Regions with a diving innervation pattern aligned most closely with sites where muscle is connected to periosteal bone by partially mineralized fiber anchors, known also as extrinsic fibers or Sharpey’s fibers^33^.

#### Type III: Meshwork

The most prevalent pattern of innervation observed consisted of a meshwork of CGRP+ sensory and TH+ sympathetic axons within the periosteum where fascia attached to the bone (Fig.2i-l). At these sites, the periosteal layer is relatively thin and the axons run close to the bone, parallel to the cortical surface. This Type III meshwork pattern is apparent in periosteal whole-mount preparations taken from regions of the bone where the absence of tendinous or extrinsic fibers allows for separation of the periosteum from adjacent tissues (Fig.2j)^7,24,34^.

### Generation of 2D and 3D maps of Type I-III periosteal innervation patterns in relation to overlying nerves and muscles

#### Nerve supply of the limb

Axons innervating the femur are primarily derived from the femoral nerve (Fig.3a). The femoral nerve runs along the anteromedial aspect of the femur and provides innervation to the muscle groups that function as knee extensors (rectus femoris and vastus muscles)^35,36^. It also provides the innervation to the anterior and medial aspects of the femur bone^37^. The obturator nerve runs nearby, also anteromedial to the femur, and innervates the hip adductors (gracilis anterior, gracilis posterior, adductor longus, adductor magnus) and some hip rotators in the gluteal region (quadratus femoris, obturator externus). There is no current evidence that the obturator innervates the femur bone. The sciatic nerve runs along the posterior aspect of the femur and innervates the posterior femoral muscles that function as hip extensors (semitendinosus, semimembranosus, caudofemoralis, biceps femoris)^35,36^. A segment of the posterior femur bone is also innervated by the sciatic nerve^38^.

**Figure 3.**
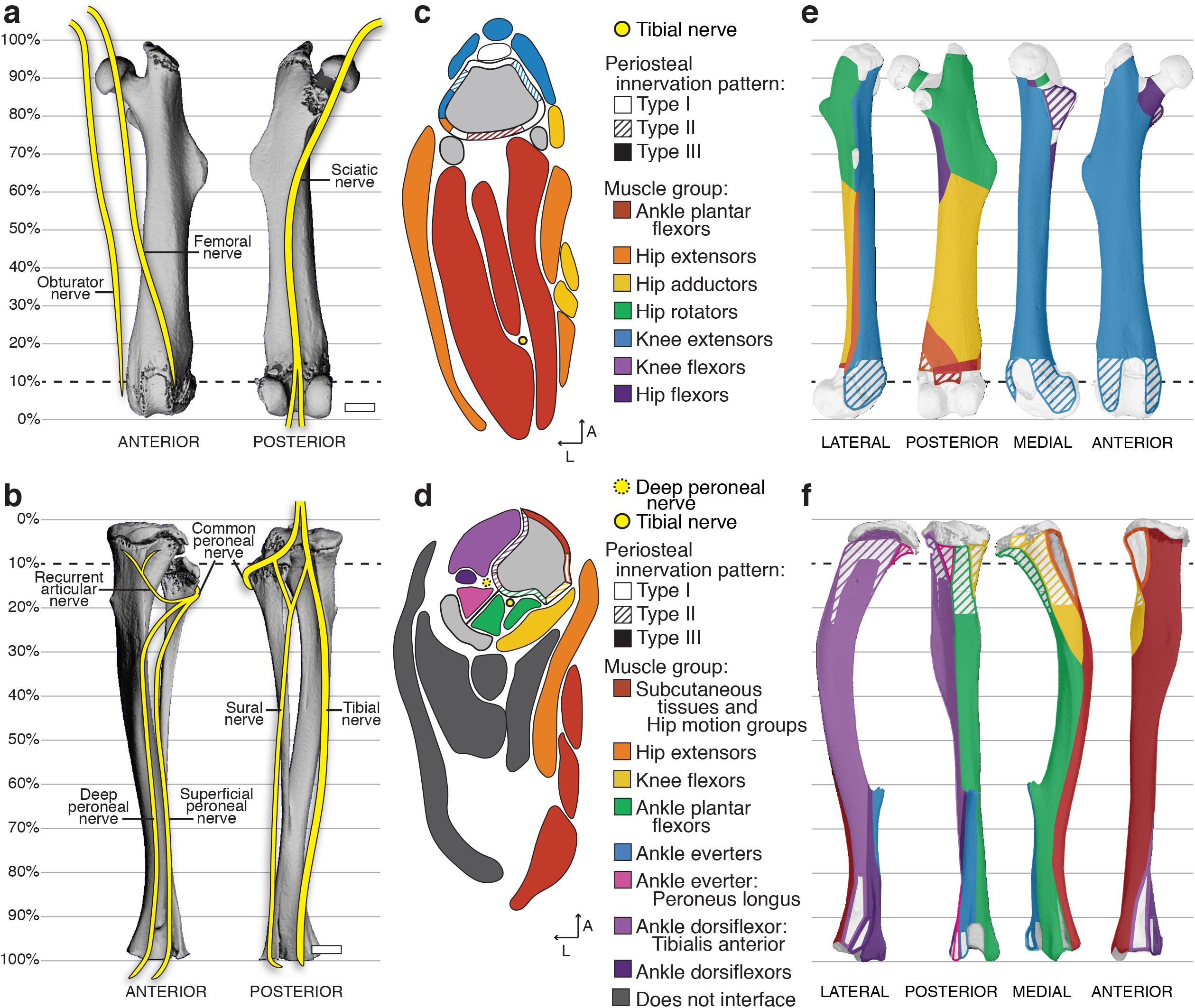
Relationships between peripheral nerves and periosteal axon patterns and the musculoskeletal system. (**a-b**) Schematic of major nerve branches in the anterior and posterior thigh (**a**) and leg (**b**) overlaid on 3D-rendered-μCT scans of the left femur and tibia-fibula, respectively. Scale bars, 1 mm. (**c,d**) Representative 2D maps of transverse cross-sections through (**c**) the thigh at the level of 10% of the total femur length proximal to the knee and (**d**) the leg at 10% of the total tibial length distal to the knee. The bone is indicated in gray and the superimposed periosteal innervation patterns are distinguished by pattern-fill and are color-coded to the interfacing muscle group indicated by the key. Detailed maps including labels of individual muscles are available in the atlases (Supp Files A-D). (**e,f**) Three-dimensional periosteal innervation maps overlaid on the lateral, posterior, medial, and anterior aspects of the μCT-rendered (**e**) femur and (**f**) tibia.

Slightly above the posterior aspect of the knee, the sciatic nerve branches into the tibial and common peroneal nerves. Innervation of the tibia bone is primarily derived from the sciatic nerve (Fig.3b) with evidence in humans for minor sensory contributions by the terminal branches of the femoral nerve in the medial regions of the proximal tibial epiphysis/metaphysis and distal epiphysis^37,38^. The tibial branch of the sciatic nerve runs down the leg on the posterior side of the tibia with the peroneal artery and veins and innervates the knee flexors, ankle plantar flexors, hip extensors, and hip motion groups in addition to the tibia bone^35,36^. By contrast, the common peroneal nerve wraps around the fibula to the front of the tibia. It sends a branch termed the recurrent articular nerve to the region of the proximal epiphysis and joint. Distally, it splits into the superficial peroneal nerve and the deep peroneal nerve. The deep peroneal nerve runs between the tibia and fibula along the interosseous membrane and, in addition to innervating the tibia bone, it sends smaller branches that innervate muscles of the ankle everter and dorsiflexor families, including the tibialis anterior^35,36^.

#### Two-dimensional muscle maps with Type I-III innervation patterns

The periosteum is a fascial connective tissue that is continuous with other layers of limb fascia including the epitenon and epimysium. To facilitate the consideration of periosteal axon patterns within the context of these boundaries, we created representative schematics showing muscle attachments and muscle labeling (Fig.3c,d). These were adapted from Charles et. al^39^ with muscle placement and innervation patterns adjusted based on our serial immunostaining analysis. Type I, II, and III innervation patterns were illustrated on the bone-tissue interface in the cross-sectional schematics and color-coded in relation to surrounding muscle attachments. This process was repeated along the length of the femur and tibia, as presented in the atlas compilation (Supplementary Files A-D) and overview section below.

#### Three-dimensional Type I-III innervation pattern maps

In addition to the 2D maps, the localization of each periosteal innervation pattern was mapped onto the surface of the femur and tibia and color-coded to match surrounding muscle groups (Fig.3e,f). Type I aneural regions were localized to the epiphyses and metaphyses of the tibia and femur. Type II regions were primarily localized in the metaphyses and along the diaphyses of the long bones. There were noted differences in the prevalence of Type II regions between B6 and C3H bones; these can be explored in more detail in the atlases, as detailed below (Supplementary Files A-D). Type III innervation was prominent on bone surfaces interfaced with loosely-adherent fascia, such as with extra-skeletal adipose tissue, epitenon, and unanchored areas of the epimysium. A Type III pattern was also found on the intraosseous fascial membranes between connected long bones, such as the tibia and fibula.

### Mapping of specialized mechanoreceptor endings

The skeleton is primarily innervated by neurons with small, finely myelinated and/or unmyelinated axons^23^. However, in specific regions, large diameter axons with specialized mechanoreceptor endings known as Pacinian corpuscles have also been identified in association with the interosseous membranes and the periosteum of large mammals^40,41^. This has not been clearly demonstrated or mapped in the mouse. To overcome this, we generated reporter mice with a floxed Ai9 allele expressing Cre recombinase under control of the *myelin protein zero (P_0_*) gene promoter to label mature myelinating Schwann cells and their precursors^42^. Schwann cells are the main glial cells of the peripheral nervous system. In addition to labeling myelinated axons, we found that the *P_0_* reporter labels the lamellar Schwann cell layers that make up the outer structure of the Pacinian corpuscle. Whole-bone tissue clearing and light-sheet microscopy of the forearm of the *P_0_*-reporter mouse revealed major nerve branches and the distribution of Pacinian corpuscles along the lateral aspect of the ulna 50-70% distal to the elbow (Fig.4a-c). In cross-sections, Pacinian corpuscles could be identified morphologically as a single myelinated sensory fiber (NF200+) in the center of a lamellar capsule that was closely approximated to, though not continuous with, the periosteum (Fig4d-f). Serial cross-section examination of Pacinian corpuscles in the limb localized these specialized mechanoreceptors exclusively to the mid-diaphyseal and distal regions of the tibia (Fig4.g-i) as mapped in the atlases (Supplementary Files A,B).

**Figure 4.**
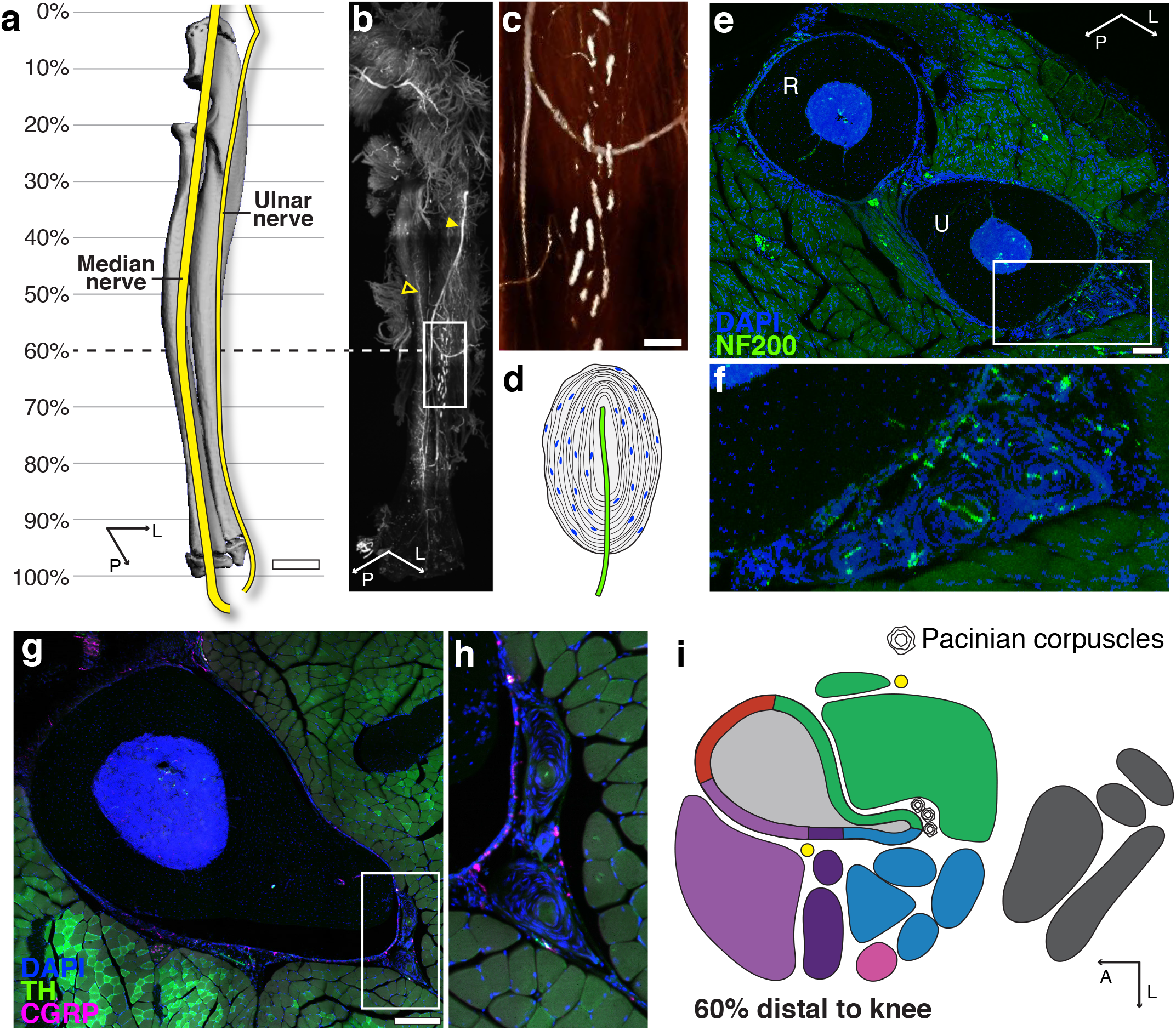
Specialized mechanoreceptor endings are associated with the skeleton of the forearm and lower leg. (**a**) Schematic of major nerve branches in the forearm overlaid on a 3D-rendered-μCT scan of the radius–ulna. Scale bar, 1 mm. (**b**) Light-sheet image of the Ai9 reporter signal in a cleared forearm from an adult Schwann cell reporter mouse (*P_0_*-cre; Ai9) revealing the ulnar nerve (filled arrowhead) and median nerve (empty arrowhead) and the distribution of Pacinian corpuscles on the ulnar surface. (**c**) High-magnification, intensity-pseudocolored image of the boxed region in b. Scale bar, 200 μm. (**d**) Drawing of a Pacinian corpuscle demonstrating the lamellar structure derived from Schwann cells surrounding a single unmyelinated ending of a large, myelinated sensory axon. (**e**) Representative confocal micrograph through a 100-μm-thick transverse cross section through the radius (R) and ulna (U) and surrounding musculature visualized by DAPI (blue) and NF200 (green) immunolabeling at 60% of the ulnar length. Scale bar, 200 μm. (**f**) High-magnification of the corpuscles on the lateral aspect of the ulna of the boxed region indicated in e. (**g**) Representative confocal projection of a 50-μm-thick transverse cross section through the leg with immunolabeling for TH+ axons (green) and CGRP+ axons (magenta) along with DAPI staining (blue) at 60% of the tibial length distal to the knee. Scale bar, 200 μm. (**h**) High-magnification of the Pacinian corpuscles on the posterior aspect of the tibia of the boxed region indicated in g. (**i**) 2D map corresponding to the confocal image in g indicating the location of Pacinian corpuscles of all strains and sexes at this level of the tibia. Additional information can be found in the supplemental tibial atlases (Supplementary Files C,D).

### Limb atlases – overview and contents

These observations and outcomes were compiled into four interactive atlases that cover the prevalence, localization, and morphology of the CGRP+ sensory and TH+ sympathetic nerve fibers in and around the femur (Supplementary Files A,B) and tibia (Supplementary Files C,D), in addition to their relationship with the overlying musculature, fascial tissue, and perilipin-positive bone marrow and extraskeletal adipocytes. Unique cross-sections were selected from serial analysis, annotated by relative distance to the knee joint. The longitudinal range of the bone that comprises similar bone morphology, as well as continuous periosteal innervation patterns and common tissue interfaces is indicated on each page to facilitate sectioning of regions of interest. Additional features and proposed use of these atlases are detailed in Figure 5.

**Figure 5.**
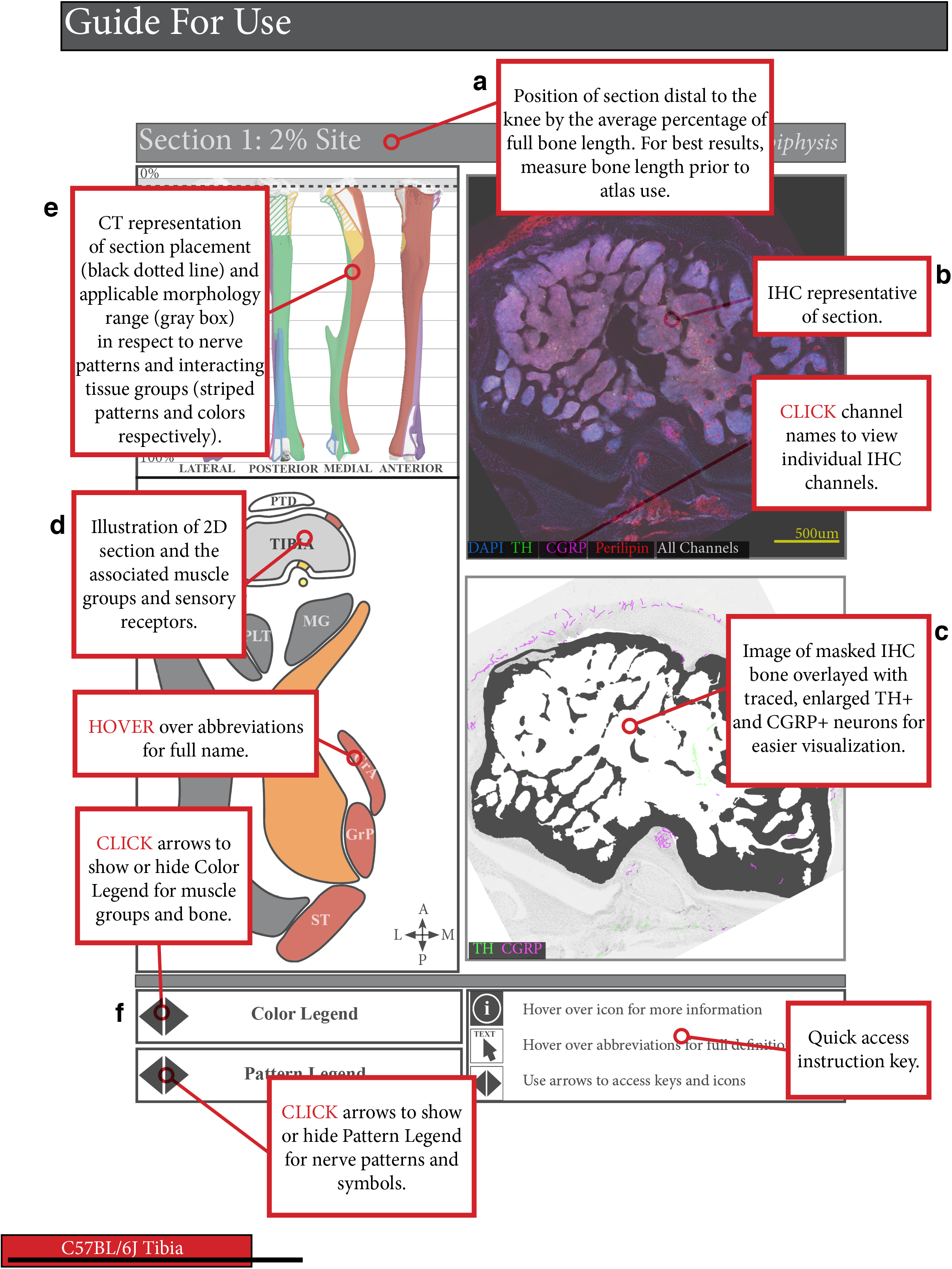
Mouse limb atlases – guide for use. **(a)** The four supplemental atlases comprise systematic analyses of serial, transverse cross-sections down the length of the femur and tibia in B6 and C3H mice, where each page details a section of the of the bone at a particular % distance from the knee. This includes (**b**) representative confocal projections of immunohistochemically labeled CGRP+ sensory and TH+ sympathetic nerve fibers, along with perilipin+ adipocytes and DAPI-stained nuclei, in and around the bone and (**c**) corresponding bone-masked, axon-traced overlays to highlight and visualize the neuroskeletal features. (**d**) In addition, the atlas contains 2D reference maps with pattern-coded periosteal innervation patterns superimposed on the bone interface and related to color-coded muscle groups and individual muscle identities. (**e**) Lastly, these 2D maps served as the basis for 3D mapping of periosteal innervation patterns and color-coded muscle groups onto the lateral, posterior, medial, and anterior aspects of the corresponding bone. (**f**) Instructions for accessing interactive features are located at the bottom of the page, such as displaying innervation pattern legend, muscle-group color legend, and full muscle identifiers, or toggling through confocal channels.

## DISCUSSION

The innervation of the skeleton is not uniform. In this manuscript, we demonstrate that peripheral axon subtype-specific densities and morphology vary by bone compartment and site and are influenced by mouse genetic strain. This dynamic, region-specific innervation of the bone and bone marrow likely influences pain and cellular function in states of skeletal homeostasis and in targeted areas of bone injury and inflammation. The maps provided here are intended to serve as a baseline reference and guide during the design and selection of regions of interest for future neuroskeletal studies in the femur, tibia, and beyond. For example, though not mapped in detail, our light sheet imaging suggests that the nerve patterns and localization of specialized mechanoreceptors within the forelimb, another common site used for skeletal research, mimic those detailed in the tibia. In addition, the tissue preparation, immunostaining, imaging, and nerve quantification methods reported here may be easily applied in any control or transgenic mouse model of targeted injury, pharmacologic intervention, or systemic disease.

Previous mapping studies in the femur (Table 1) showed that the periosteum is the most densely innervated compartment, where CGRP+ peptidergic sensory axons predominate, while TH+ sympathetic axons are more prevalent in the marrow cavity. For the first time, we corroborate these findings in the tibia, and additionally report a strain-dependent innervation density and CGRP:TH ratio, and a strain-independent proximal-to-distal increase in bone marrow axon density. Neural contributions to adaptive loading, as well as fracture pain and healing, have been largely ascribed to local sensory fibers^17,43,44^, which predominate in the periosteum. Neural contributions to hematopoiesis and skeletal metastasis have been attributed to sympathetic activity^45,46^, in line with the intimate association of sympathetic fibers with the arteriolar vasculature and their relative abundance in the marrow. Both arms of the peripheral nervous system have been implicated in skeletal homeostasis, suggesting counter-regulatory mechanisms afforded by sensory and sympathetic innervation co-exist to maintain bone homeostasis, for example by coordinating opposing effects on vascular tone or osteoblast function^1^. By contrast, non-overlapping sensory and sympathetic axons within the bone marrow may be suggestive of spatially-defined mechanisms regulating unique progenitor cell pools^18,20^.

As detailed throughout the atlases, bone is intimately associated with the nervous, vascular, and muscular systems. Periosteal axon patterning, which we have characterized and defined as Type I-III, is aligned to connections of muscle and fascia to the bone. The endocortical surface, by contrast, is relatively aneural. Axonal orientation is less defined in the marrow, but TH+ fibers generally cluster around central arterioles, while CGRP+ axons are more disperse (Supplementary Files A-D). Particularly exciting potential exists in overlaying this neuroskeletal map with whole-bone maps of load-induced strain and adaptation^47^, vascularity^48^, and stem-cell distribution^49,50^. For example, strain maps could be related to periosteal innervation patterns to give insight to their functional role in load-induced adaptation; Type II axons may integrate muscle cues to the loaded bone while axons in Type III regions might amplify bone formation on surfaces that lack large connective tissue strain. Additionally, this work highlights the need to consider the role of Schwann cells in the bidirectional cross-talk between nerves and bone. Schwann cells not only insulate axons, but can also initiate and modulate neural signaling, as well as contribute to tissue repair^51–53^, and remain an underexplored component of the neuroskeletal system.

As we move forward, it is clear that a systematic and comprehensive map of skeletal innervation related to region-specific bone surfaces and bone marrow compartments is fundamental to assigning physiologic functions to local neuronal populations. In addition, the neuroskeletal field needs standardized nomenclature to facilitate more comprehensive and cohesive studies. In this resource, we have provided protocols and easy-to-apply methods for axon visualization and quantification. In addition, we have presented four interactive atlases that can be applied as a reference during study design, implementation, and interpretation. Lastly, based on this work, we propose that future manuscripts report several basic metrics to help to unify the field. Specifically, (#1) section orientation (transverse, longitudinal), (#2) the analyzed level along the length of the bone *(e.g.* 40% from the knee), (#3) the bone surface (*e.g.* anterior tibial ridge), and (#4) the axon pattern(s) analyzed (*e.g.* Types II and/or III). This will help to ensure reproducibility between studies while moving the field toward development of new therapies for treatment of musculoskeletal pain and disease.

## ACKNOWLEDGEMENTS

This work was supported by grants from the National Institutes of Health including U01-DK116317 (E.L.S.) and T32-AR060719 (J.M.B.). We are thankful to the Musculoskeletal Research Center (MRC, P30-AR074992) and the Washington University Center for Cellular Imaging (WUCCI), with particular thanks to James Fitzpatrick and Peter Bayguinov, for their support. In addition, we would like to thank Jeff Stirman at SmartSPIM for generating the lightsheet image used in this publication.

## AUTHOR CONTRIBUTIONS (CRediT taxonomy)

Conceptualization: M.R.L, J.M.B., A.B., E.L.S.

Data curation: M.R.L, J.M.B., A.B., I.S., E.L.S.

Formal analysis: A.B., I.S., E.L.S.

Funding acquisition: E.L.S.

Investigation: M.R.L, J.M.B., A.B., I.S., E.L.S.

Methodology: M.R.L, J.M.B., A.B., E.L.S.

Project administration: E.L.S.

Resources: E.L.S.

Software: M.R.L, J.M.B., A.B., E.L.S.

Supervision: E.L.S.

Validation: M.R.L, J.M.B., A.B., E.L.S.

Visualization: M.R.L, J.M.B., A.B., E.L.S.

Writing – original draft: M.R.L, J.M.B., A.B., E.L.S.

Writing – review & editing: M.R.L, J.M.B., A.B., I.S., E.L.S.

## METHODS

### Table of key resources

2 Table of key resources
| REAGENT or RESOURCE | SOURCE | IDENTIFIER |
| --- | --- | --- |
| MOUSE MODELS |  |  |
| C57BL6J | Jackson Laboratories | Strain 000664 |
| C3H/HeJ | Jackson Laboratories | Strain 000659 |
| C57BL6J: Schwann Cell Reporter Mouse | Dr. Jeffrey Milbrandt | P<sub>0</sub>-Cre (JAX Strain 017927) and Ai9-flox/flox (JAX Strain 007909) |
| CHEMICALS |  |  |
| 10% neutral buffered formalin | Fisher Scientific | 23-245684 |
| EDTA | Sigma-Aldrich | E5134 |
| DAPI | Sigma-Aldrich | D9542 |
| OCT mounting media | Fisher HealthCare | 23-730-571 |
| Triton X-100 | Sigma-Aldrich | 9002-93-1 |
| Fluoromount-G | Thermo Fisher Scientific | 00-4958-02 |
| Donkey serum | Sigma-Aldrich | D9663 |
| Acrylamide/bis-acrylamide, 40% solution | Sigma-Aldrich | A7802 |
| Paraformaldehyde | Electron Microscopy Sciences | 15710 |
| Histodenz | Sigma Aldrich | D2158-100G |
| NNN'N'-Tetrakis(2-Hydroxypropyl)ethylenediamine-Quadrol | Sigma Aldrich | 122262-1L |
| VA-044 | Wako | 27776-21-2 |
| SDS Micropellets | Thermo Fisher Scientific | BP8200-500 |
| Antibodies for immunostaining and western blot | Detailed in Table S1 |  |
| SOFTWARE |  |  |
| GraphPad Prism | GraphPad | v8.4.3 |
| SCANCO Medical microCT systems | Scanco Medical AG |  |
| FIJI | ImageJ |  |
| Microsoft Excel | Microsoft |  |
| Adobe Acrobat Pro DC | Adobe Creative Cloud |  |
| Adobe InDesign | Adobe Creative Cloud |  |
| Adobe Illustrator | Adobe Creative Cloud |  |
| Adobe Photoshop | Adobe Creative Cloud |  |
| ORS Dragonfly | Object Research Systems |  |
| IMAGING SYSTEMS |  |  |
| Scanco μCT 40 | Scanco Medical AG | N/A |
| Spinning Disk Confocal Microscope | Nikon CSU-X1 | N/A |
| Smart SPIM Lightsheet Imaging System | Life Canvas Technologies |  |
| Leica Confocal Microscope | Leica TCS SPEII confocal |  |
| OTHER |  |  |
| PicoLab Rodent Diet 20 | LabDiet | 5053 |
| Colorfrost Plus Microscope Slides | Fisher Scientific | 12-550-18 |
| Cryostat | Leica Biosystems |  |

### Mice

All work was performed as approved by the animal use and care committee at Washington University (Saint Louis, MO, USA). Mice were housed on a 12-hour light/dark cycle and fed *ad libitum* (PicoLab 5053, LabDiet). C57BL6J (JAX Strain #000664) and C3H/HeJ (JAX Strain #000659) mice were obtained from Jackson Laboratories. Schwann cell reporter mice were a kind gift of Dr. Jeffrey Milbrandt and were generated by crossing the following strains: *P_0_*-Cre (JAX Strain 017927)^42^ and Ai9-flox/flox (JAX Strain 007909)^54^.

### Histology and Immunostaining

#### Tissue isolation and embedding

Mice were anesthetized with Ketamine/Xylazine and perfused through the left ventricle of the heart with 10 mL phosphate buffered saline followed by 10 mL 10% neutral buffered formalin (NBF, Fisher Scientific 23-245684). For all experiments, collected tissues were post-fixed in 10% NBF for 24-hours. After fixation, tissues were washed for 2 h in diH_2_O. Bones were fully decalcified in 14% EDTA (Sigma-Aldrich E5134), pH 7.4 and equilibrated in 30% sucrose solution prior to embedding in OCT mounting media (Fisher HealthCare 23-730-571).

#### Frozen immunostaining and imaging

Embedded tissues were cut at 50 μm on a cryostat (Leica). For immunostaining, cut sections on Colorfrost Plus glass slides (Fisher Scientific 12-550-18) were blocked in 10% donkey serum in TNT buffer prior to incubation for 48-hours with primary antibodies at 4°C (Table S1). After washing 3x 5 min, secondary antibodies in TNT buffer were applied for 24 h at 4°C (Table S1). The sections were then washed 3x 5 min in TNT buffer, incubated in DAPI (Sigma-Aldrich) for 5 min, and washed again prior to mounting with Fluoromount-G (Thermo Fisher Scientific 00-4958-02). For Fig.1,2 and Supplemental Files A-D, serial tiled images were taken with a 10x objective on a Nikon spinning disk confocal microscope (μm/px = 0.650, step size = 2.5 μm Number of steps = 21). Stitching of the nd2 multipoint files was done using the FIJI Stitching plugin “snake by rows”. For Fig.5, serial tiles images were obtained using a Leica confocal at 10x (μm/px = 0.6968, step size = 2.5 μm, number of steps= 41) with automatic stitching through the LASX Software.

**Table S1.** Antibodies used for immunostaining of fixed frozen tissue.

| Table S1. Antibodies used for immunostaining of fixed frozen tissue. |  |  |  |  |
| --- | --- | --- | --- | --- |
| Primary Antibody (Vendor, Cat. No) | Dilution | Secondary Antibody | Fluorophore | Dilution |
| Anti-Calcitonin Gene Related Peptide (Bio-rad, 1720-9007) | 1:1000 | Donkey Anti-Goat (Jackson ImmunoResearch, 705-606-147) | AlexaFluor-647 | 1:500 |
| Anti-Tyrosine Hydroxylase (Abcam, UK, ab152) | 1:1000 | Donkey Anti-Rabbit (Jackson ImmunoResearch, 711-546-152) | AlexaFluor-488 | 1:500 |
| Anti-Perilipin (Progen Biotechnik, Germany, GP29) | 1:500 | Donkey Anti-Guinea Pig (Jackson ImmunoResearch, 706-165-148) | Cy3 | 1:500 |
| Neurofilament 200 (NF200) (Millipore, AB5539) | 1:5000 | Donkey Anti-Chicken (Jackson ImmunoResearch, 703-545-155) | AlexaFluor-488 | 1:500 |

### Confocal Image Segmentation and Analysis

#### Selection of quantification regions

From the longitudinal array of slides created for each bone, four sections were chosen corresponding to the metaphysis, mid-diaphysis with tibial ridge, diaphysis proximal to the TFJ, and diaphysis just distal to the TFJ. For each of the above regions, sections were chosen based on morphological similarity, including bone shape and attachment sites.

#### Generation of tissue masks

A combined cancellous and cortical bone mask was generated simultaneously by thresholding max projections of FITC channel (green) in FIJI^55^. Optimal upper and lower thresholds were chosen to remove muscle, connective tissue, marrow and other background cellularity (some speckle and small holes in bone were allowed at this stage). After thresholding, bone masks were manually corrected based on the source file in ImageJ by clearing non-bone elements using the freehand selection tool. A median filter (radius = 11) was applied to smooth the mask. Medullary masks were generated by inverting the bone mask in ImageJ, drawing a circle inside the cortical ring using the freehand selection tool, and clearing outside the selection. Cancellous and cortical bone masks were produced using the medullary mask, clearing inside or outside the marrow cavity. Lastly holes in the cancellous and cortical bone masks were filled using the FIJI binary operation Fill Holes. In cases where cortical bone completely enclosed the marrow cavity a small path was cleared from the cortex prior to performing the binary operation, and subsequently repaired.

The periosteum was defined as a thin, densely cellular layer adjacent to cortical bone. Based on the max projections of the FITC channel in FIJI, upper and lower thresholds were chosen to segment the periosteum. Using previous masks, the bone and marrow were deleted from the periosteal mask. Masks were manually corrected in ImageJ using the freehand selection tool. The DAPI channel was frequently referenced to differentiate periosteum from other connective tissues and fascia. A median filter (radius = 6) was applied to smooth the mask and lastly holes in the periosteum were filled using the FIJI binary operation Fill Holes to create a continuous layer. In cases where the periosteum made a complete ring a small path was cleared from the periosteum prior to performing the binary operation, and subsequently repaired.

#### Cortical and cancellous BV/TV calculation

The area of the cortical bone, cancellous bone, and marrow cavity were calculated in FIJI from the masks created by thresholding-based segmentation described above. Cortical BV/TV was calculated as cortical bone area/(cortical bone area + marrow area) and cancellous BV/TV was calculated as cancellous bone area/marrow area.

#### Axon tracing and quantification

TH+ and CGRP+ axons were independently traced with the Simple Neurite Tracer plugin to obtain total axon length. The volume of the periosteum and marrow cavity were calculated as the area multiplied by a section thickness of 50 μm. Volumetric density of fiber lengths was calculated as the total length of fibers traced divided by the corresponding volume for the region traced. Identification of nerves within the bone marrow is more challenging than at other sites due to non-specific autofluorescence, poor light penetration, and the presence of antibody-dependent cellular staining. This can result in staining artifacts, as noted with an information icon (“i”) when they appear in the atlas (Supp Files A-D). Strategies to overcome these limitations include technical optimization, tissue clearing, tiled confocal images of thick frozen sections to capture large volume datasets, and strict morphologic criteria for nerve selection.

#### Axon tracing image overlays

TH+ and CGRP+ axon tracings were exported from the Simple Neurite Tracer plugin in FIJI^56^. This exported file preserves the coordinates of the traced axons and assigns each traced segment a line with a width of one pixel. Exported nerve masks were dilated four times using the binary processing tool, increasing the width of each traced axon to 9 pixels for the purposes of visualization. After this, bone and nerve masks were imported into Adobe Photoshop and overlaid on a grayscale max projection of the FITC and DAPI channels from the original image with cortical bone and bone marrow removed. This results in a visually accurate map of axon length, branching, orientation and relative density. However, features such as absolute nerve diameter are not preserved.

### MicroCT and Maps

Representative two and three-dimensional maps of the Type I, II, and III periosteal innervation patterns were generated after comprehensive review of serial images generated every 150 μm along the length of the femur and tibia from four independent mice (12-week old B6 male, B6 female, C3H male, C3H female). A representative 3D model, color-coded to match surrounding functional muscle groups, was then reconstructed in Adobe Photoshop using scans of the tibiae of 12-week-old B6 and C3H male mice performed on a Scanco μCT 40 (Scanco Medical AG, Bassersdorf, Switzerland; 55 kVp, 145 μA, 300 ms integration time, 16 μm voxel size). 2D Schematics showing muscle attachments and muscle labeling were generated in Adobe Illustrator. These were adapted from Charles et. al^39^ with muscle placement and innervation patterns adjusted based on our serial section analysis. Type I, II, and III innervation patterns were illustrated on the bone-tissue interface in the 2D schematics and color-coded in relation to surrounding muscle attachments and fascial structures. The final atlas was compiled with Adobe InDesign.

### Whole Mount Tissue Clearing and Imaging

Tissue clearing was performed as previously described with minor modification^57^. Briefly, mice were intracardially perfused with 20 mL 0.02M PBS followed by 20 mL of fresh 4% paraformaldehyde. Immediately after perfusion, dissected tissues were post-fixed in 4% PFA at 4° overnight. The post-fixed limb was then incubated in A4P0 hydrogel overnight (5 mL 40% acrylamide, 45 mL of PBS, and 0.125 g of VA044) and polymerized at 38° for 6 hours. The forelimb was then incubated in 8% SDS at 37° for 5 days to fully clear muscle and bone tissue. After 2 days of thorough PBS washing, the forelimb was incubated in 25% quadrol at 37° for 2 days. After another 2 days of thorough washing in PBS, the bones were incubated in 1.42 and 1.48 RIMS of Histodenz for one day each before imaging. Images were taken with 4x objective on the SmartSPIM imaging system through the full length of the bone and 3D reconstructed with ORS Dragonfly Software for visualization of the P_0_+ Schwann cells.

### Statistics

Statistical analyses were performed in GraphPad Prism. Specific tests are indicated in the figure legends. A p-value of less than 0.050 was considered statistically significant.

## DATA AVAILABILITY

All relevant data are available from the authors upon reasonable request. Raw images will be made available on the SPARC data portal (https://sparc.science/) – to be released with final publication.

## REFERENCES

1. Brazill, J. M., Beeve, A. T., Craft, C. S., Ivanusic, J. J. & Scheller, E. L. Nerves in Bone: Evolving Concepts in Pain and Anabolism. J. Bone Miner. Res. 34, 1393–1406 (2019).

2. O’Neill, T. W. & Felson, D. T. Mechanisms of Osteoarthritis (OA) Pain. Curr. Osteoporos. Rep. 16, 611–616 (2018).

3. Ivanusic, J. J. Molecular Mechanisms That Contribute to Bone Marrow Pain. Front. Neurol. 8, 458 (2017).

4. Mantyh, P. W. The neurobiology of skeletal pain. Eur. J. Neurosci. 39, 508–519 (2014).

5. Sayilekshmy, M. et al. Innervation is higher above Bone Remodeling Surfaces and in Cortical Pores in Human Bone: Lessons from patients with primary hyperparathyroidism. Sci. Rep. 9, (2019).

6. Mach, D. B. et al. Origins of skeletal pain: sensory and sympathetic innervation of the mouse femur. Neuroscience 113, 155–166 (2002).

7. Martin, C. D., Jimenez-Andrade, J. M., Ghilardi, J. R. & Mantyh, P. W. Organization of a unique net-like meshwork of CGRP+ sensory fibers in the mouse periosteum: implications for the generation and maintenance of bone fracture pain. Neurosci. Lett. 427, 148–152 (2007).

8. Fukuta, H., Mitsui, R., Takano, H. & Hashitani, H. Contractile properties of periosteal arterioles in the guinea-pig tibia. Pflugers Arch. 469, 1203–1213 (2017).

9. Fukuta, H., Mitsui, R., Takano, H. & Hashitani, H. Neural regulation of the contractility of nutrient artery in the guinea pig tibia. Pflugers Arch. 472, 481–494 (2020).

10. Wee, N. K. Y., Lorenz, M. R., Bekirov, Y., Jacquin, M. F. & Scheller, E. L. Shared Autonomic Pathways Connect Bone Marrow and Peripheral Adipose Tissues Across the Central Neuraxis. Front. Endocrinol. 10, (2019).

11. Ferrell, W. R., Khoshbaten, A. & Angerson, W. J. Responses of bone and joint blood vessels in cats and rabbits to electrical stimulation of nerves supplying the knee. J. Physiol. 431, 677–687 (1990).

12. Tomlinson, R. E. et al. NGF-TrkA Signaling by Sensory Nerves Coordinates the Vascularization and Ossification of Developing Endochondral Bone. Cell Rep. 16, 2723–2735 (2016).

13. Heffner, M. A., Genetos, D. C. & Christiansen, B. A. Bone adaptation to mechanical loading in a mouse model of reduced peripheral sensory nerve function. PloS One 12, e0187354 (2017).

14. Sample, S. J. et al. Functional adaptation to loading of a single bone is neuronally regulated and involves multiple bones. J. Bone Miner. Res. 23, 1372–1381 (2008).

15. Madsen, J. E. et al. Fracture healing and callus innervation after peripheral nerve resection in rats. Clin. Orthop. 230–240 (1998).

16. Zhang, Y. et al. Implant-derived magnesium induces local neuronal production of CGRP to improve bone-fracture healing in rats. Nat. Med. 22, 1160–1169 (2016).

17. Chartier, S. R. et al. Exuberant sprouting of sensory and sympathetic nerve fibers in nonhealed bone fractures and the generation and maintenance of chronic skeletal pain. Pain 155, 2323–2336 (2014).

18. Hu, B. et al. Sensory nerves regulate mesenchymal stromal cell lineage commitment by tuning sympathetic tones. J. Clin. Invest. 130, 3483–3498 (2020).

19. Zhu, S. et al. Subchondral bone osteoclasts induce sensory innervation and osteoarthritis pain. J. Clin. Invest. 129, 1076–1093 (2019).

20. Katayama, Y. et al. Signals from the sympathetic nervous system regulate hematopoietic stem cell egress from bone marrow. Cell 124, 407–421 (2006).

21. Imai, S., Tokunaga, Y., Maeda, T., Kikkawa, M. & Hukuda, S. Calcitonin gene-related peptide, substance P, and tyrosine hydroxylase-immunoreactive innervation of rat bone marrows: an immunohistochemical and ultrastructural investigation on possible efferent and afferent mechanisms. J. Orthop. Res. 15, 133–140 (1997).

22. Robles, H. et al. Characterization of the bone marrow adipocyte niche with three-dimensional electron microscopy. Bone (2018).

23. Ivanusic, J. J. Size, neurochemistry, and segmental distribution of sensory neurons innervating the rat tibia. J. Comp. Neurol. 517, 276–283 (2009).

24. Castañeda-Corral, G. et al. The majority of myelinated and unmyelinated sensory nerve fibers that innervate bone express the tropomyosin receptor kinase A. Neuroscience 178, 196–207 (2011).

25. Chartier, S. R., Mitchell, S. A. T., Majuta, L. A. & Mantyh, P. W. The Changing Sensory and Sympathetic Innervation of the Young, Adult and Aging Mouse Femur. Neuroscience 387, 178–190 (2018).

26. Judex, S., Garman, R., Squire, M., Donahue, L.-R. & Rubin, C. Genetically based influences on the site-specific regulation of trabecular and cortical bone morphology. J. Bone Miner. Res. 19, 600–606 (2004).

27. Beamer, W. G., Donahue, L. R., Rosen, C. J. & Baylink, D. J. Genetic variability in adult bone density among inbred strains of mice. Bone 18, 397–403 (1996).

28. Scheller, E. L. et al. Region-specific variation in the properties of skeletal adipocytes reveals regulated and constitutive marrow adipose tissues. Nat. Commun. 6, 7808 (2015).

29. Orwoll, E. S. Toward an expanded understanding of the role of the periosteum in skeletal health. J. Bone Miner. Res. 18, 949–954 (2003).

30. Duchamp de Lageneste, O. et al. Periosteum contains skeletal stem cells with high bone regenerative potential controlled by Periostin. Nat. Commun. 9, 773 (2018).

31. Apostolakos, J. et al. The enthesis: a review of the tendon-to-bone insertion. Muscles Ligaments Tendons J. 4, 333–342 (2014).

32. Suzuki, D., Murakami, G. & Minoura, N. Histology of the bone-tendon interfaces of limb muscles in lizards. Ann. Anat. Anat. Anz. 184, 363–377 (2002).

33. Aaron, J. E. Periosteal Sharpey’s fibers: a novel bone matrix regulatory system? Front. Endocrinol. 3, (2012).

34. Thai, J., Kyloh, M., Travis, L., Spencer, N. J. & Ivanusic, J. J. Identifying spinal afferent (sensory) nerve endings that innervate the marrow cavity and periosteum using anterograde tracing. J. Comp. Neurol. (2020) doi:10.1002/cne.24862.

35. Greene, E. C. Anatomy of the rat. (Hafner Pub. Co., 1955).

36. Waxman, S. G. Clinical neuroanatomy. (McGraw-Hill Education/Medical, 2013).

37. Ultrasound-Guided Femoral Nerve Block. NYSORA https://www.nysora.com/techniques/lower-extremity/ultrasound-guided-femoral-nerve-block/ (2018).

38. Ultrasound-Guided Sciatic Nerve Block. NYSORA https://www.nysora.com/regional-anesthesia-for-specific-surgical-procedures/lower-extremity-regional-anesthesia-for-specific-surgical-procedures/foot-and-anckle/ultrasound-guided-sciatic-nerve-block-2/ (2018).

39. Charles, J. P., Cappellari, O., Spence, A. J., Hutchinson, J. R. & Wells, D. J. Musculoskeletal Geometry, Muscle Architecture and Functional Specialisations of the Mouse Hindlimb. PloS One 11, e0147669 (2016).

40. Ferrington, D. G., Rowe, M. J. & Tarvin, R. P. Actions of single sensory fibres on cat dorsal column nuclei neurones: vibratory signalling in a one-to-one linkage. J. Physiol. 386, 293–309 (1987).

41. Zelená, J. Survival of Pacinian corpuscles after denervation in adult rats. Cell Tissue Res. 224, 673–683 (1982).

42. Feltri, M. L. et al. P0-Cre transgenic mice for inactivation of adhesion molecules in Schwann cells. Ann. N. Y. Acad. Sci. 883, 116–123 (1999).

43. Li, Z. et al. Fracture repair requires TrkA signaling by skeletal sensory nerves. J. Clin. Invest. 129, 5137–5150 (2019).

44. Tomlinson, R. E. et al. NGF-TrkA signaling in sensory nerves is required for skeletal adaptation to mechanical loads in mice. Proc. Natl. Acad. Sci. U. S. A. 114, E3632–E3641 (2017).

45. Maryanovich, M. et al. Adrenergic nerve degeneration in bone marrow drives aging of the hematopoietic stem cell niche. Nat. Med. 24, 782–791 (2018).

46. Clément-Demange, L., Mulcrone, P. L., Tabarestani, T. Q., Sterling, J. A. & Elefteriou, F. β2ARs stimulation in osteoblasts promotes breast cancer cell adhesion to bone marrow endothelial cells in an IL-1β and selectin-dependent manner. J. Bone Oncol. 13, 1–10 (2018).

47. Carriero, A. et al. Spatial relationship between bone formation and mechanical stimulus within cortical bone: Combining 3D fluorochrome mapping and poroelastic finite element modelling. Bone Rep. 8, 72–80 (2018).

48. Grüneboom, A. et al. A network of trans-cortical capillaries as mainstay for blood circulation in long bones. Nat. Metab. 1, 236–250 (2019).

49. Acar, M. et al. Deep imaging of bone marrow shows non-dividing stem cells are mainly perisinusoidal. Nature 526, 126–130 (2015).

50. Kokkaliaris, K. et al. Adult blood stem cell localization reflects the abundance of reported bone marrow niche cell types and their combinations. Blood. [published online ahead of print]. (2020).

51. Yamazaki, S. et al. Nonmyelinating Schwann cells maintain hematopoietic stem cell hibernation in the bone marrow niche. Cell 147, 1146–1158 (2011).

52. Stierli, S., Imperatore, V. & Lloyd, A. C. Schwann cell plasticity-roles in tissue homeostasis, regeneration, and disease. Glia 67, 2203–2215 (2019).

53. Abdo, H. et al. Specialized cutaneous Schwann cells initiate pain sensation. Science 365, 695–699 (2019).

54. Madisen, L. et al. A robust and high-throughput Cre reporting and characterization system for the whole mouse brain. Nat. Neurosci. 13, 133–140 (2010).

55. Schindelin, J. et al. Fiji: an open-source platform for biological-image analysis. Nat. Methods 9, 676–682 (2012).

56. Longair, M. H., Baker, D. A. & Armstrong, J. D. Simple Neurite Tracer: open source software for reconstruction, visualization and analysis of neuronal processes. Bioinforma. Oxf. Engl. 27, 2453–2454 (2011).

57. Greenbaum, A. et al. Bone CLARITY: Clearing, imaging, and computational analysis of osteoprogenitors within intact bone marrow. Sci. Transl. Med. 9, (2017).

