## Supplementary File C for "A neuroskeletal atlas of the mouse limb"

### C57BL/6J Tibia

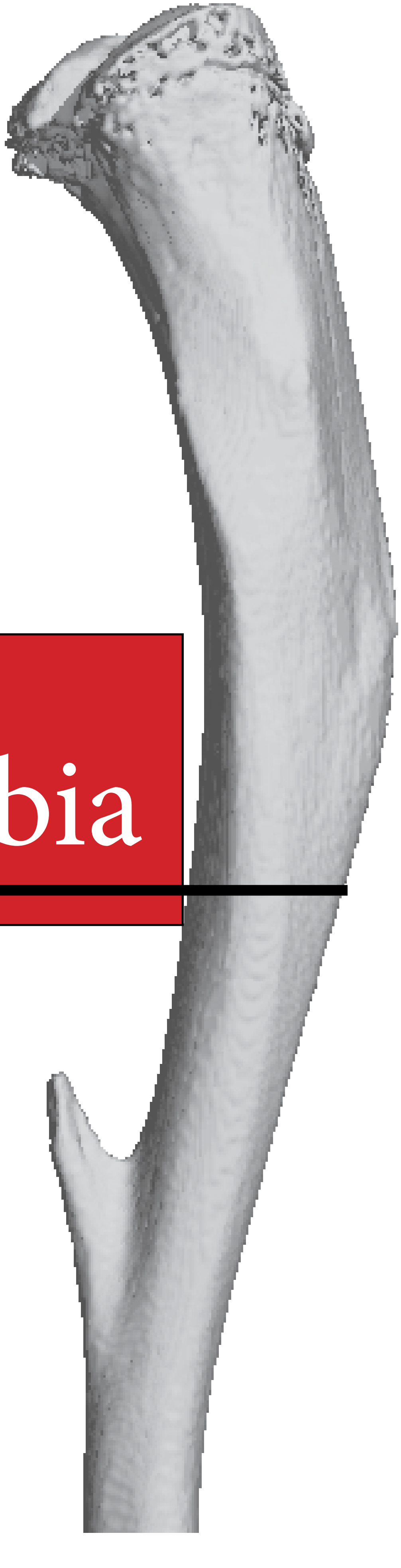

#### Methods

The following atlas was constructed from 50 µm frozen serial sections through the tibiae of five male and female C57BL/6J 12-week-old mice. Calcitonin gene-related peptide (CGRP)-positive peptidergic sensory fibers [Bio-rad, 1720-9007] and tyrosine hydroxylase (TH)-positive sympathetic fibers [Abcam, AB152], as well as perilipin-positive adipocytes [Progen, GP29] were labeled by immunohistochemistry and detected with fluorescent-conjugated secondary antibodies along with DAPI-stained nuclei [Sigma, D9542]. These sections were imaged and analyzed using a Nikon CSU-X1 spinning disk confocal system. Cancellous and cortical bone masks were generated by thresholding max projections in ImageJ. TH+ and CGRP+ axons were traced with the Simple Neurite Tracer plugin. Bone masks and axon traces were overlaid on a grayscale max projection of the FITC and DAPI channels. 3D maps were generated using a 3D model reconstructed from scans of the tibiae of 12-week-old male mice performed on a Scanco µCT 40. 2D schematics showing muscle attachments and muscle labeling were adapted from Charles et. al (1) with muscle placement adjusted based on our serial section analysis. Innervation patterns were illustrated on the bone-tissue interface in our 2D schematics in relation to surrounding muscle attachments and fascial structures. Pacinian corpuscles, specialized nerve endings responsive to mechanical distortion and vibration, were identified by morphology and are also annotated throughout. The final atlas was compiled with Adobe InDesign.

#### Notes and Limitations

Representative IHC images in the following pages include both male (sections 1-9) and female (sections 10-11) mice; please note that the bone morphology and innervation pattern are relatively conserved, while peripheral fat content is increased in females. As with all immunostaining procedures, staining and sectioning artifacts are possible during processing. These are noted throughout the atlas when present. Click the information buttons for more details about a specific artifact. Nerve tracings rely on the relative strength of immunolabeling and assessment of axon morphology. Axons parallel to the section plane were positively stained, curvilinear structures of 0.2-5 µm in diameter; those perpendicular to the section plane were small punctate structures, also of 1-5 µm diameter, that continued through the entire depth of the 50 µm z-stack. Axons oriented longitudinally, perpendicular to the plane of section, will appear smaller than their actual length when visualized in 2D.

#### Acknowledgements

Special thanks to the Washington University Center for Cellular Imaging and the Washington University Musculoskeletal Research Center for imaging equipment and technical support.

#### Bibliography

1. Charles JP, Cappellari O, Spence AJ, Hutchinson JR, Wells DJ. Musculoskeletal geometry, muscle architecture and functional specialisations of the mouse hindlimb. PLoS One. 2016 Apr 26;11(4):e0147669.

### Guide For Use

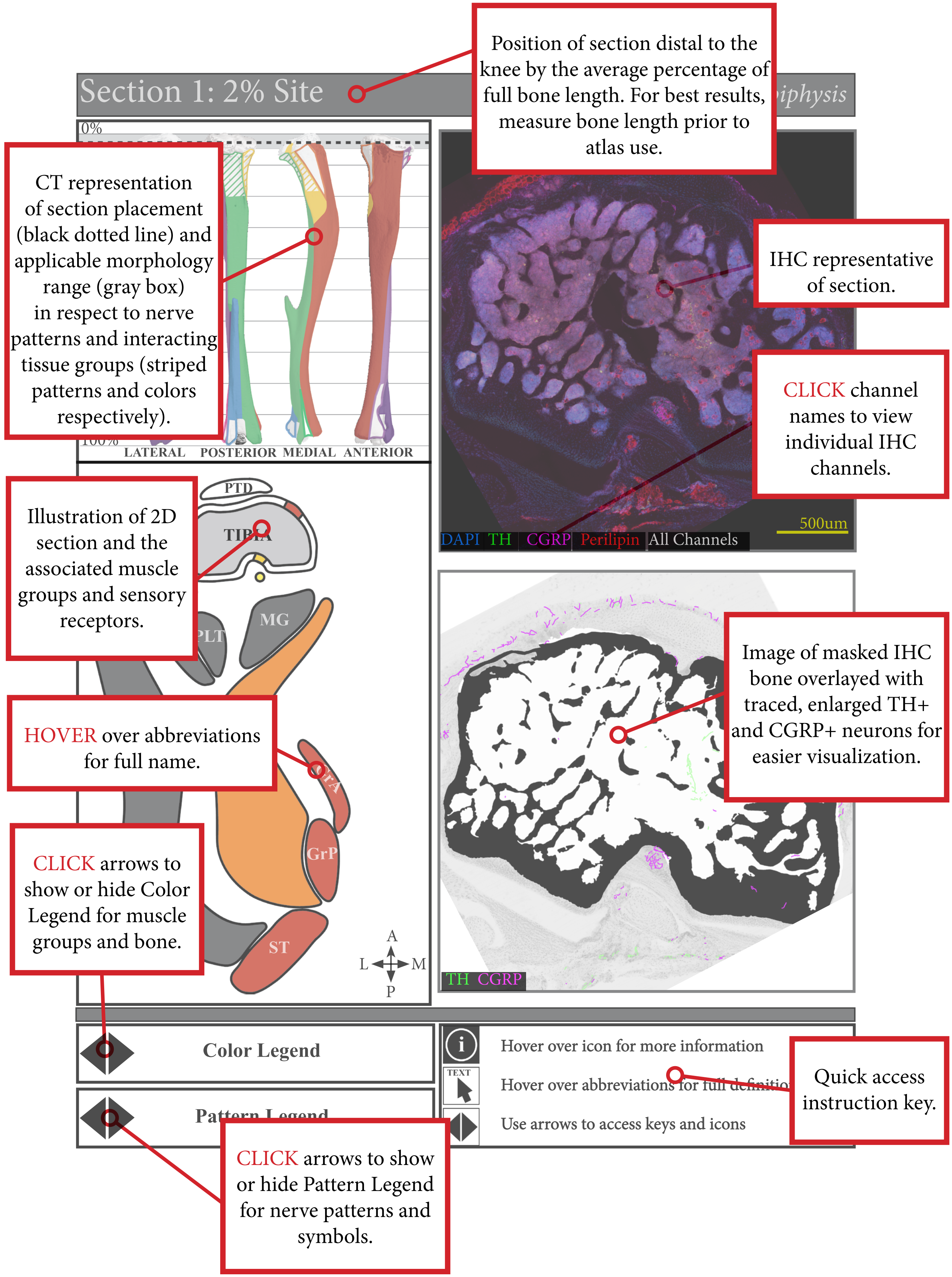

### Legend

##### Muscle Groups

- Subcutaneous Tissues and Hip Motion Groups
- Hip Extensors
- Knee Flexors
- Ankle Plantar Flexors
- Ankle Everters
- Ankle Everter: Peroneus Longus
- Ankle Dorsiflexor: Tibialis Anterior
- Ankle Dorsiflexors
- Do not interact with studied bone surfaces

##### Bones

- Tibia, Fibula, and Foot

##### Innervation Patterns

- Type I: Aneural
- Type II: Anchored
- Type III: Fascial/ Parallel

##### Icons

- Pacinian Corpuscles
- Major Nerve Branches

### Abbreviations

| | | | |
|---|---|---|---|
| **BF** | Bicep femoralis posterior | **PDQ** | Peroneus digiti quinti/quarti |
| **CGRP** | Calcitonin gene-related peptide | **PL** | Peroneus longus |
| **DAPI** | 4',6-diamidino-2-phenylindole | **PLT** | Plantaris |
| **EDL** | Extensor digitorum longus | **PT** | Peroneus tertius |
| **EHL** | Extensor hallucis longus | **PTD** | Patellar tendon |
| **F** | Fibula | **POP** | Popliteus |
| **FDL** | Flexor digitorum longus | **SM** | Semimembranosus |
| **GrA** | Gracilis anterior | **SOL** | Soleus |
| **GrP** | Gracilis posterior | **ST** | Semitendinosus |
| **LG** | Lateral gastrocnemius | **TA** | Tibialis anterior |
| **MG** | Medial gastrocnemius | **TH** | Tyrosine hydroxylase |
| **PB** | Peroneus brevis | **TP** | Tibialis posterior |

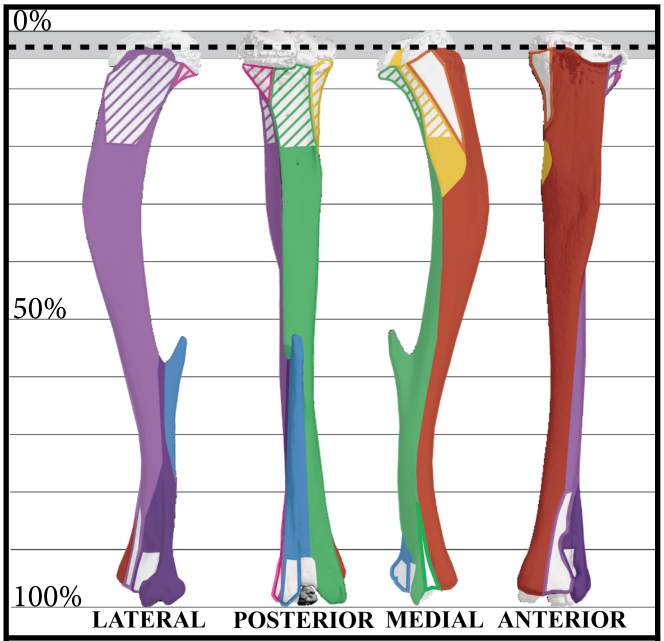


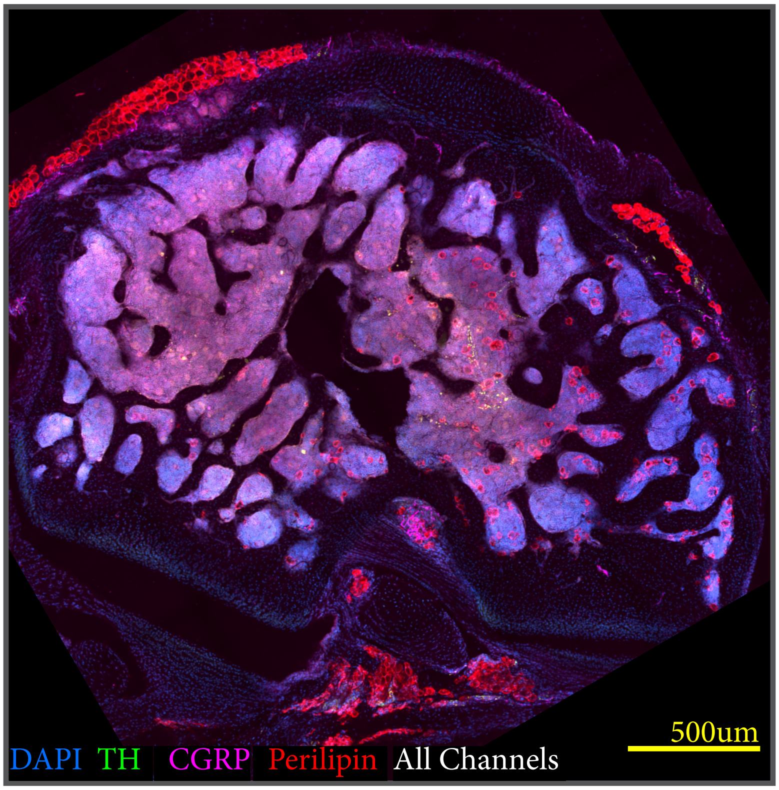


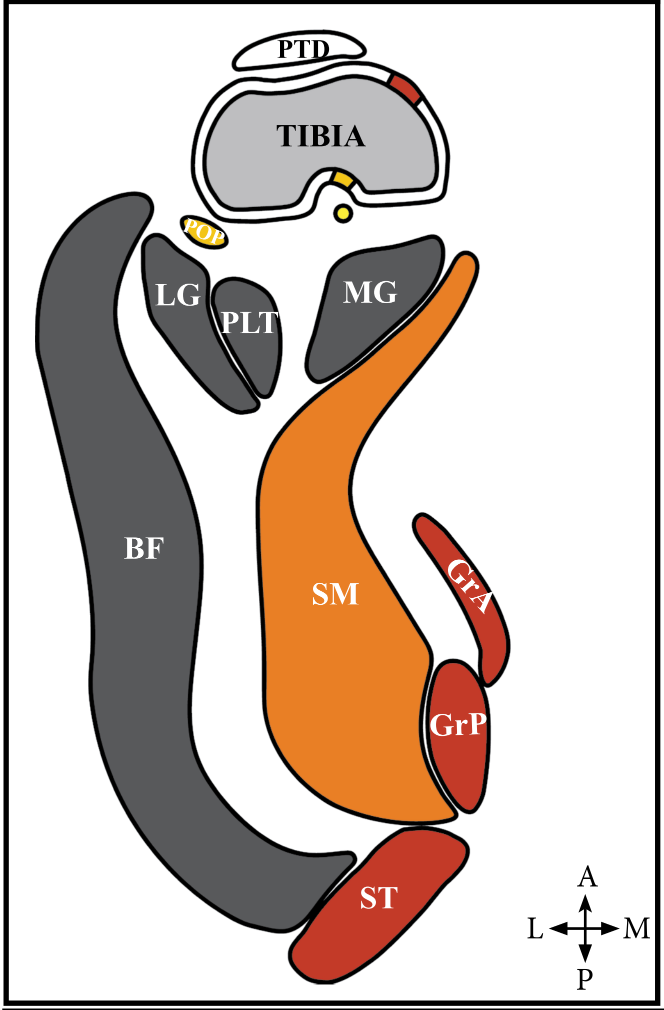


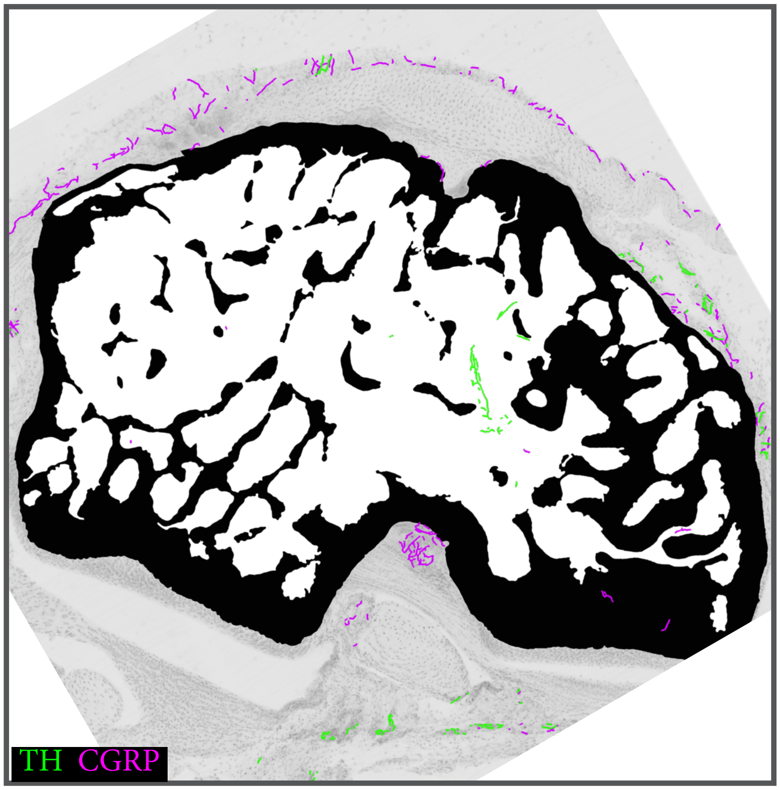


**Color Legend**

**Pattern Legend**

Hover over icon for more information

TEXT Hover over abbreviations for full definition

Use arrows to access keys and icons

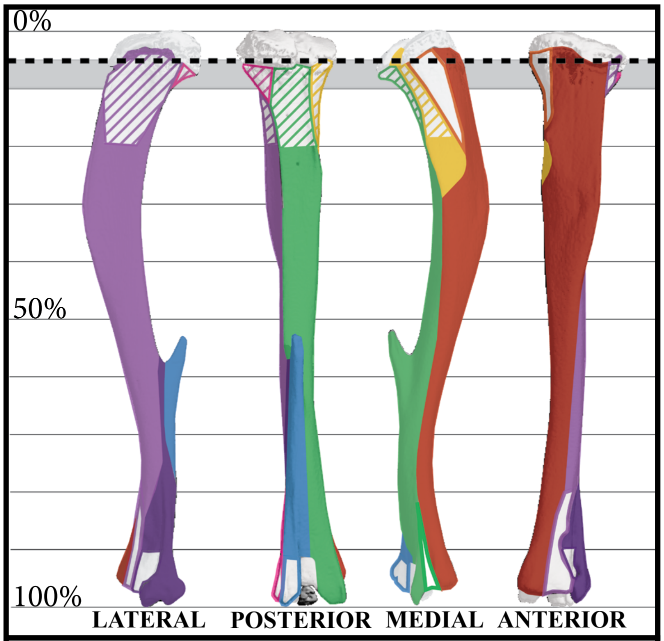


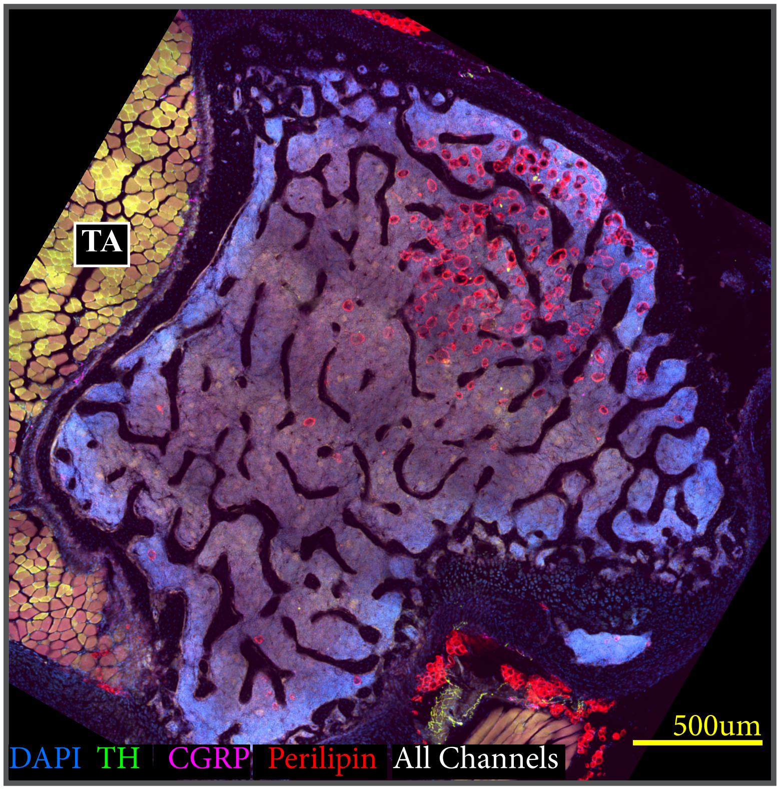


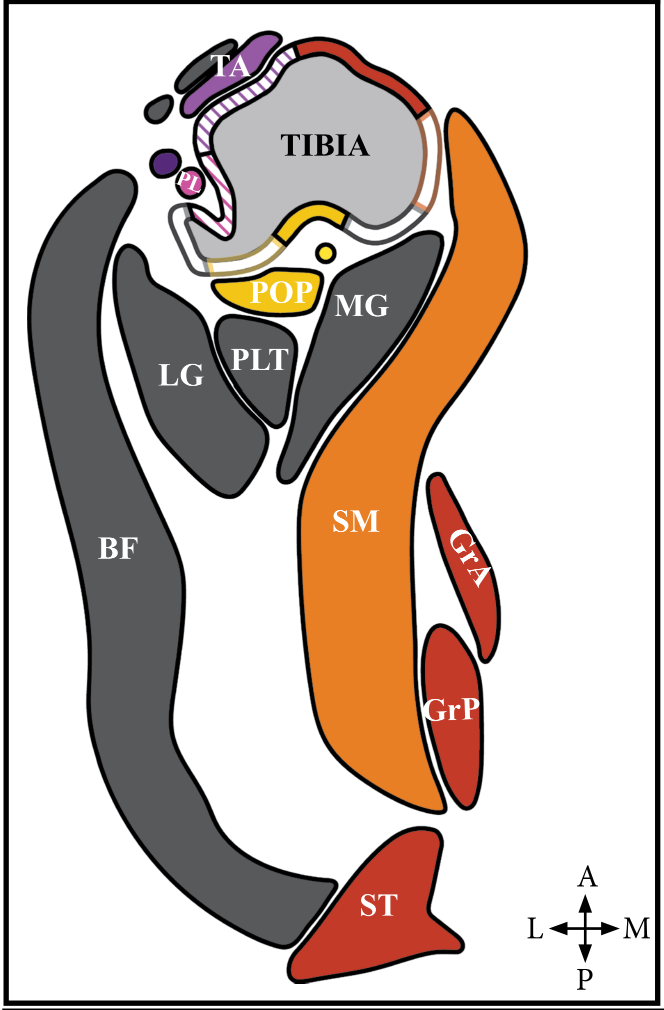


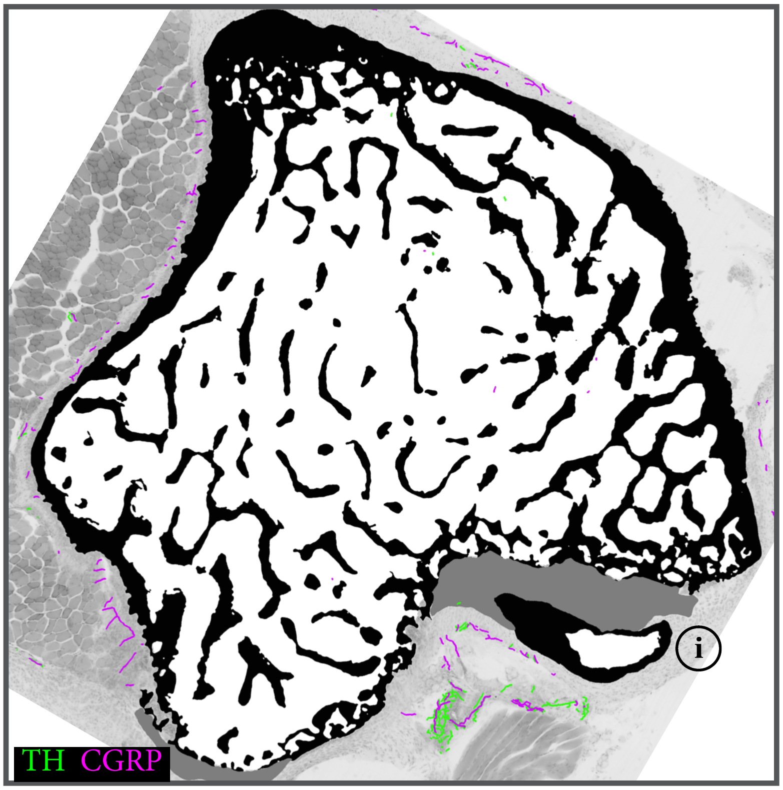


**Color Legend**

**Pattern Legend**

Hover over icon for more information

Hover over abbreviations for full definition

Use arrows to access keys and icons

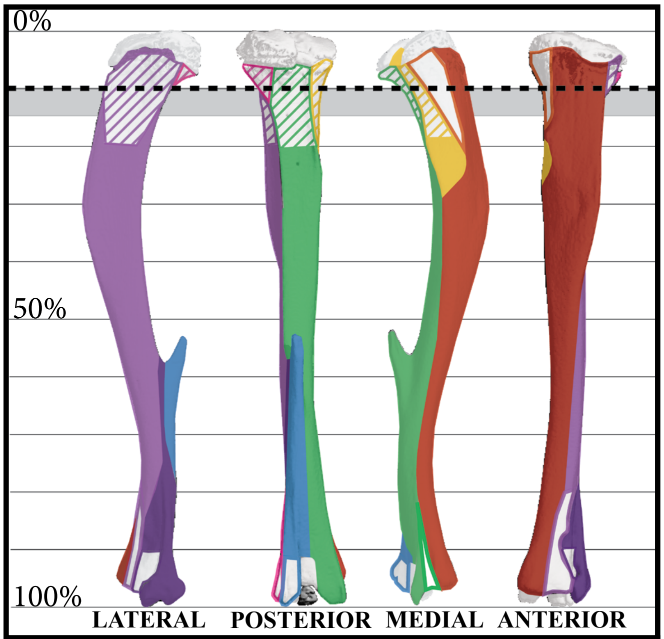


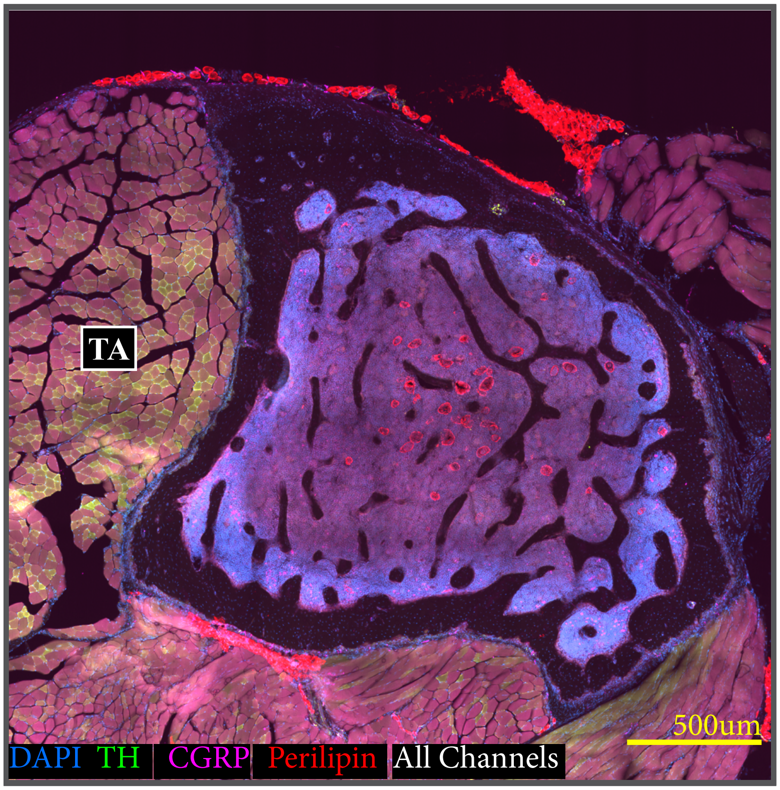


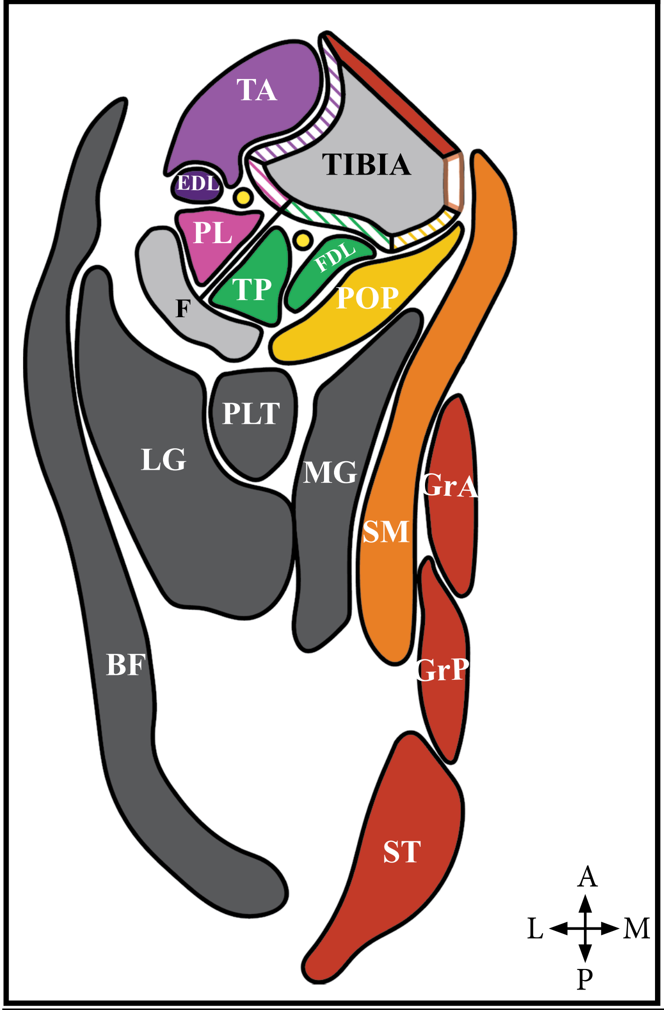


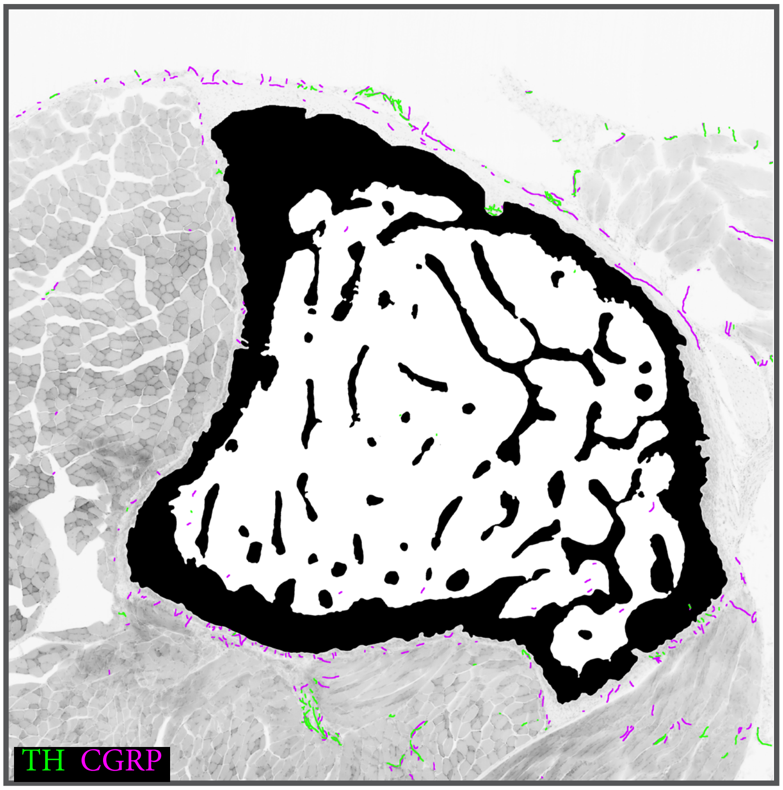


**Color Legend**

**Pattern Legend**

Hover over icon for more information

TEXT Hover over abbreviations for full definition

Use arrows to access keys and icons

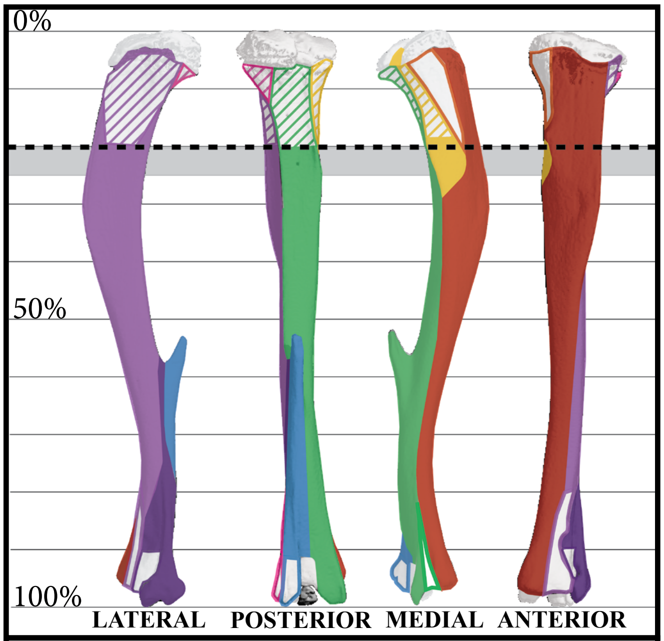


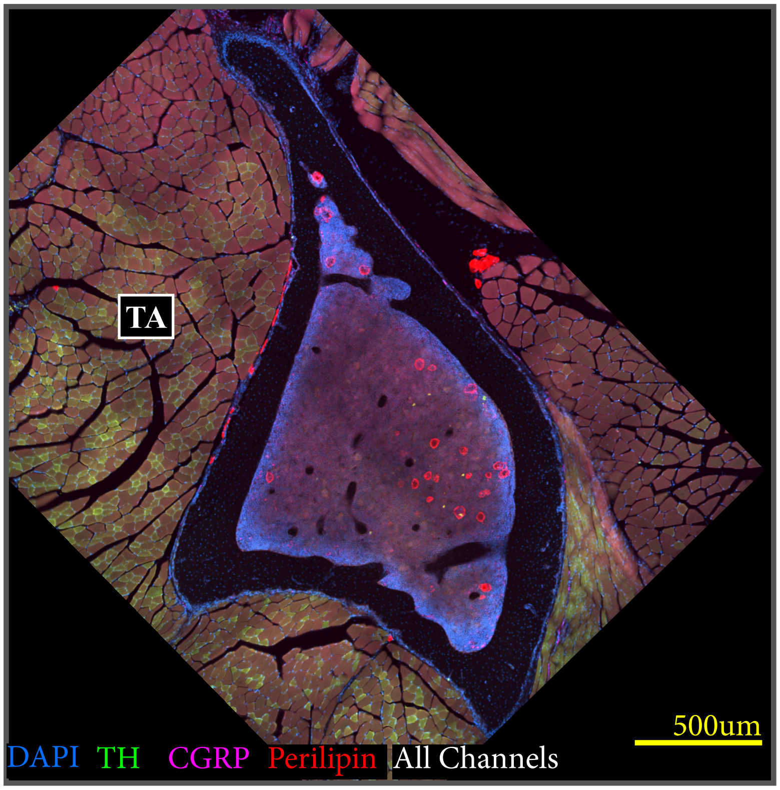


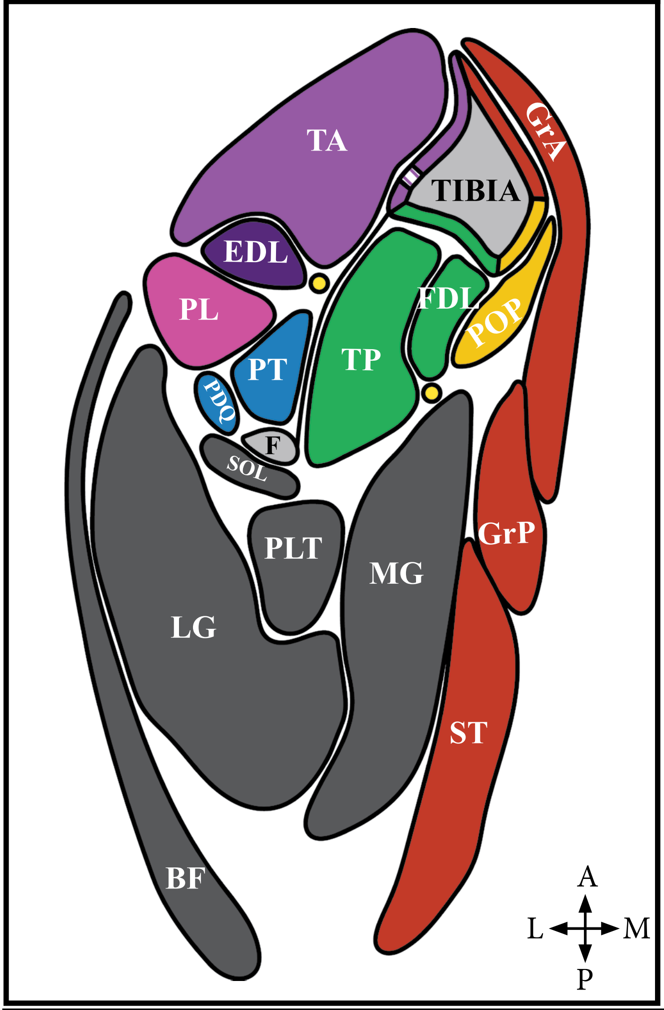


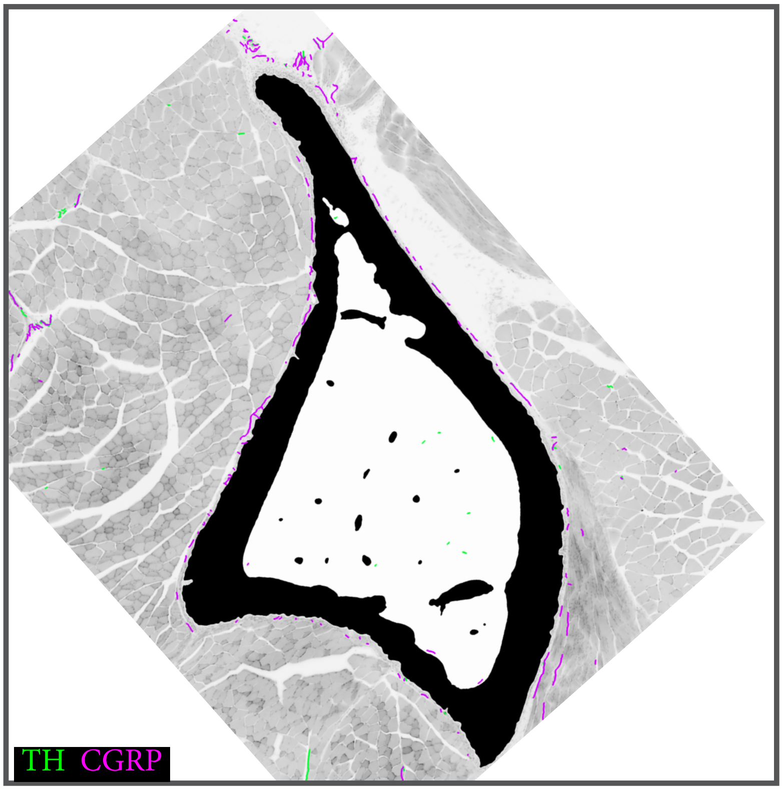


**Color Legend**

**Pattern Legend**

Hover over icon for more information

Hover over abbreviations for full definition

Use arrows to access keys and icons

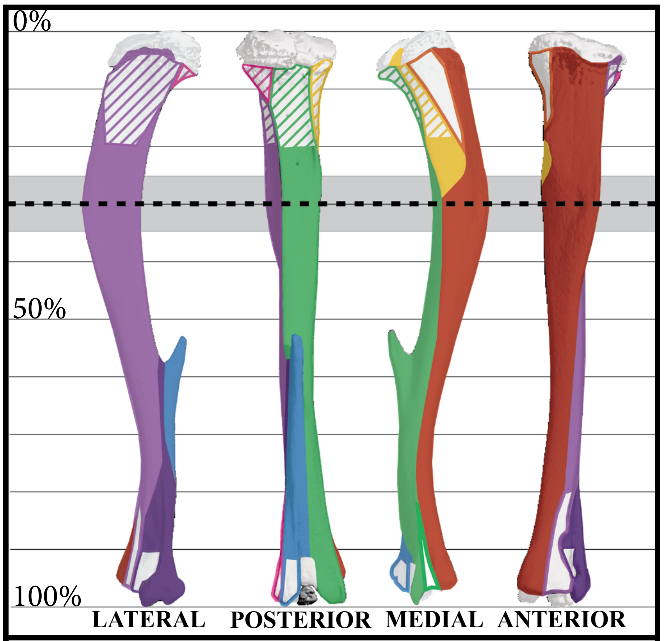


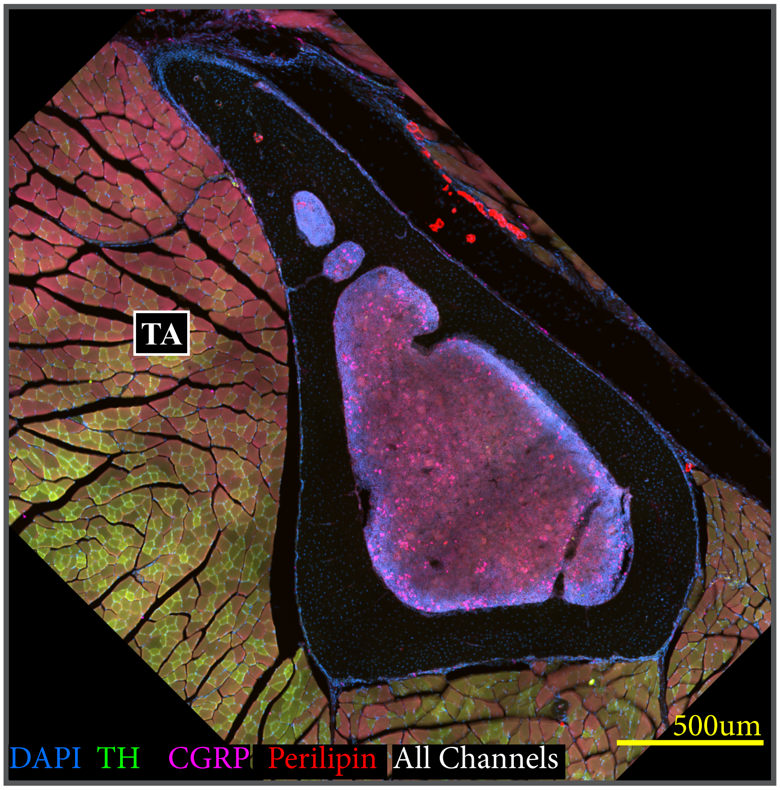


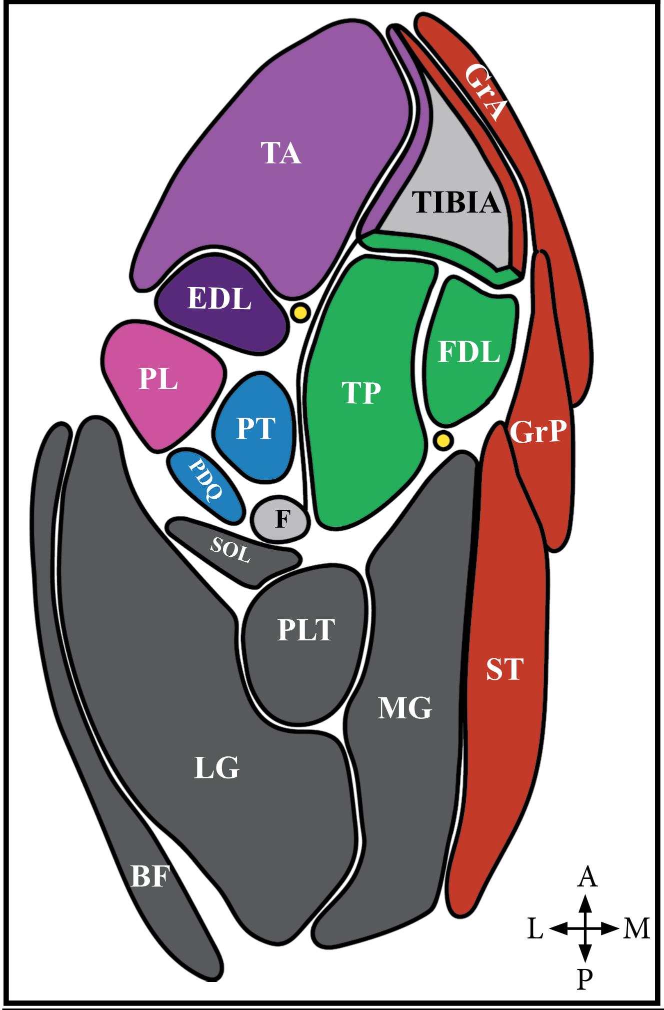


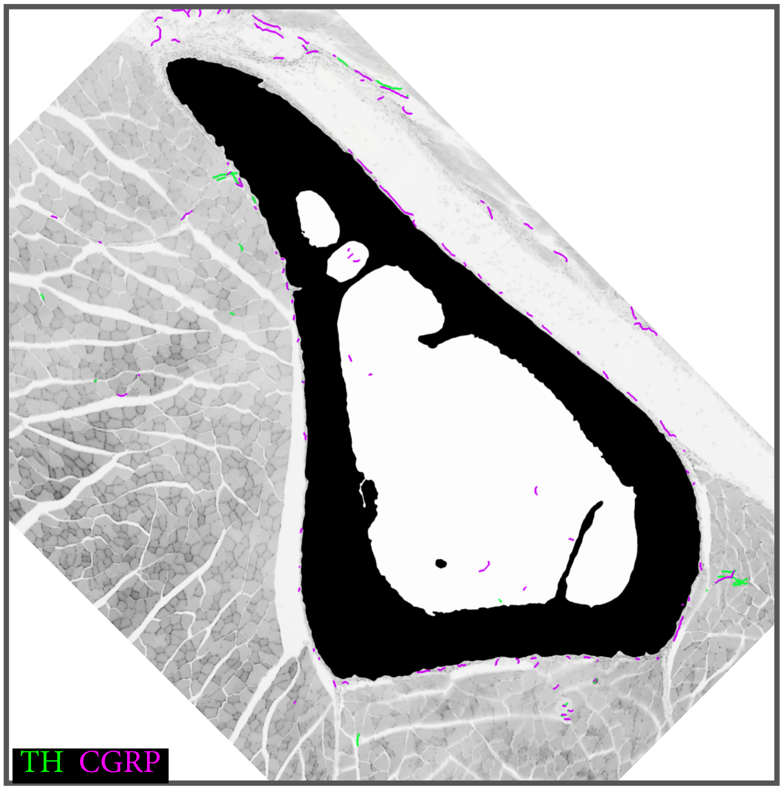


Color Legend

Pattern Legend

Hover over icon for more information

Hover over abbreviations for full definition

Use arrows to access keys and icons

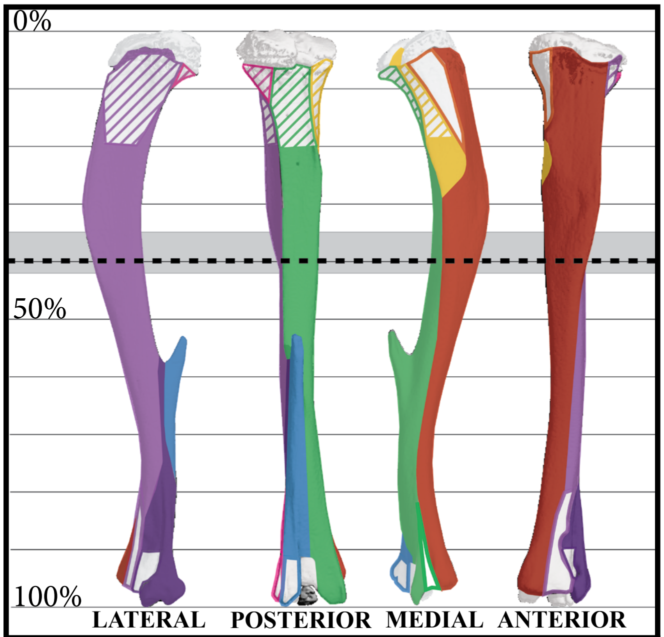


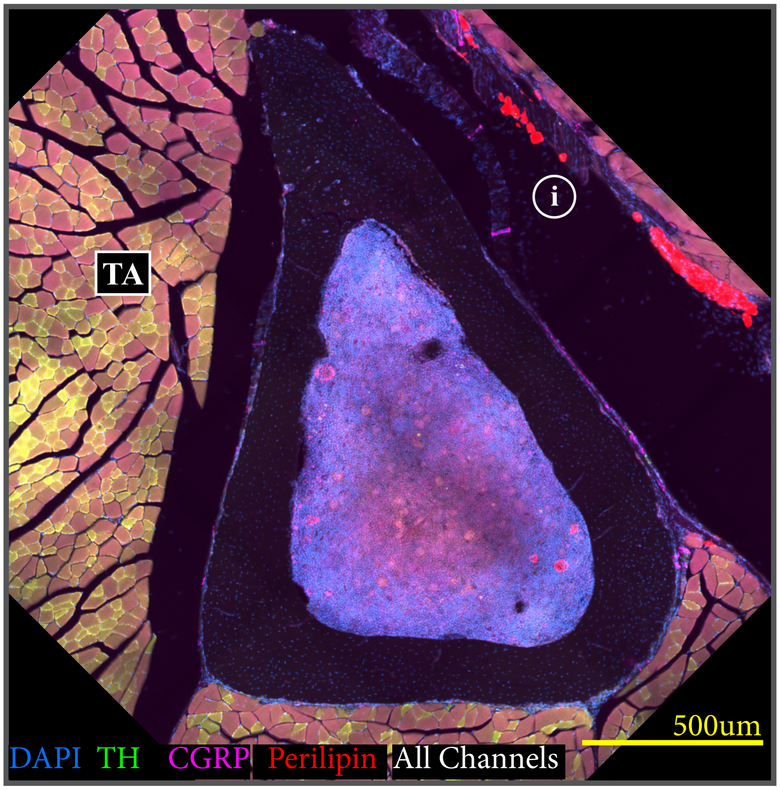


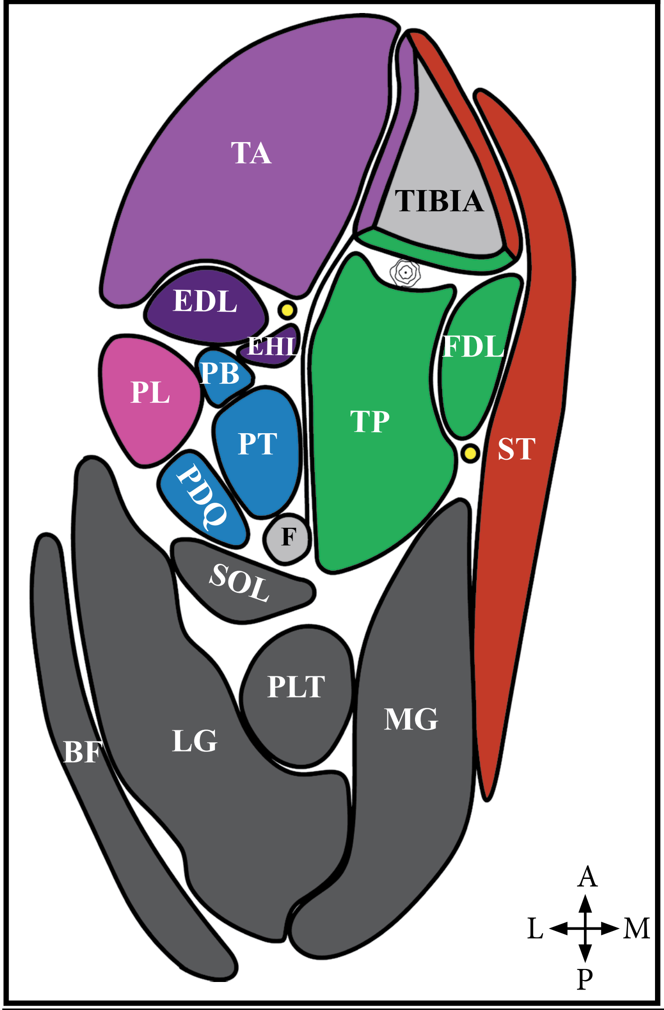


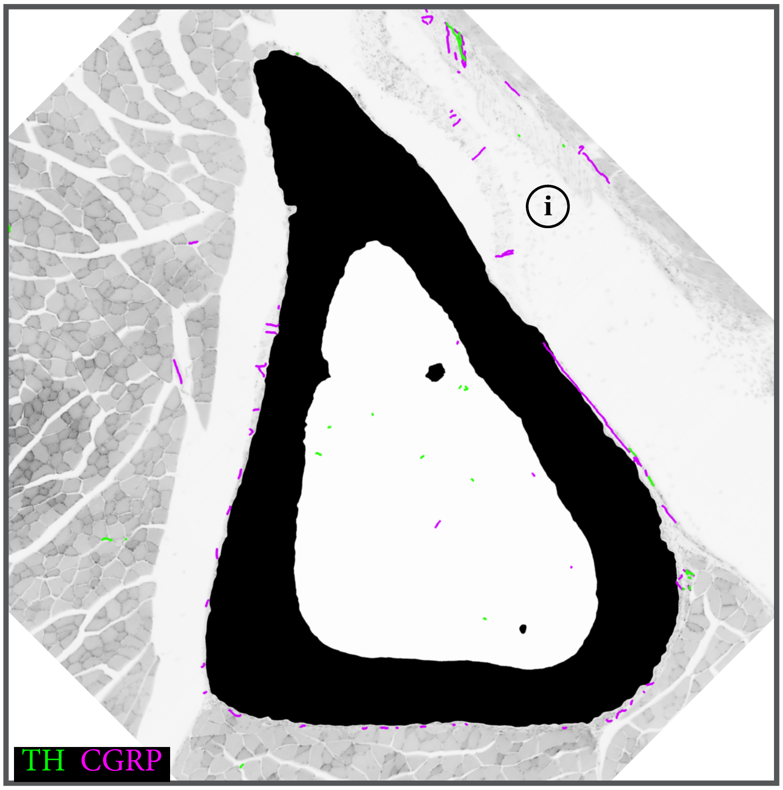


**Color Legend**

**Pattern Legend**

Hover over icon for more information

Hover over abbreviations for full definition

Use arrows to access keys and icons

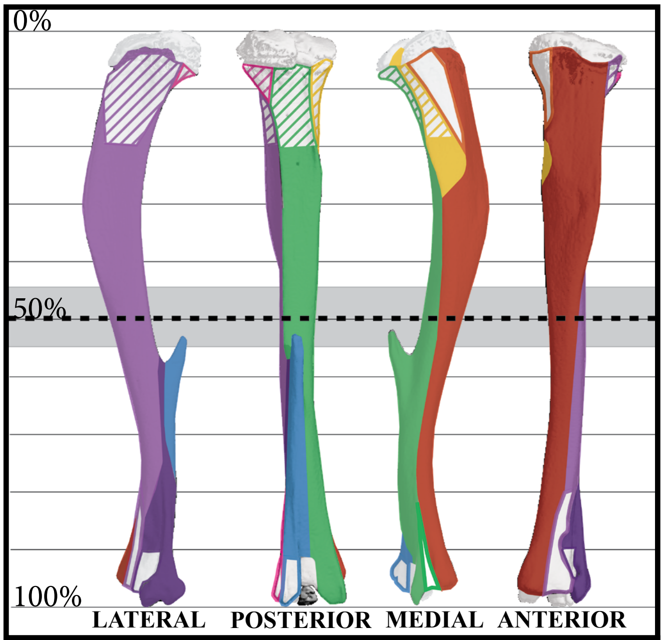


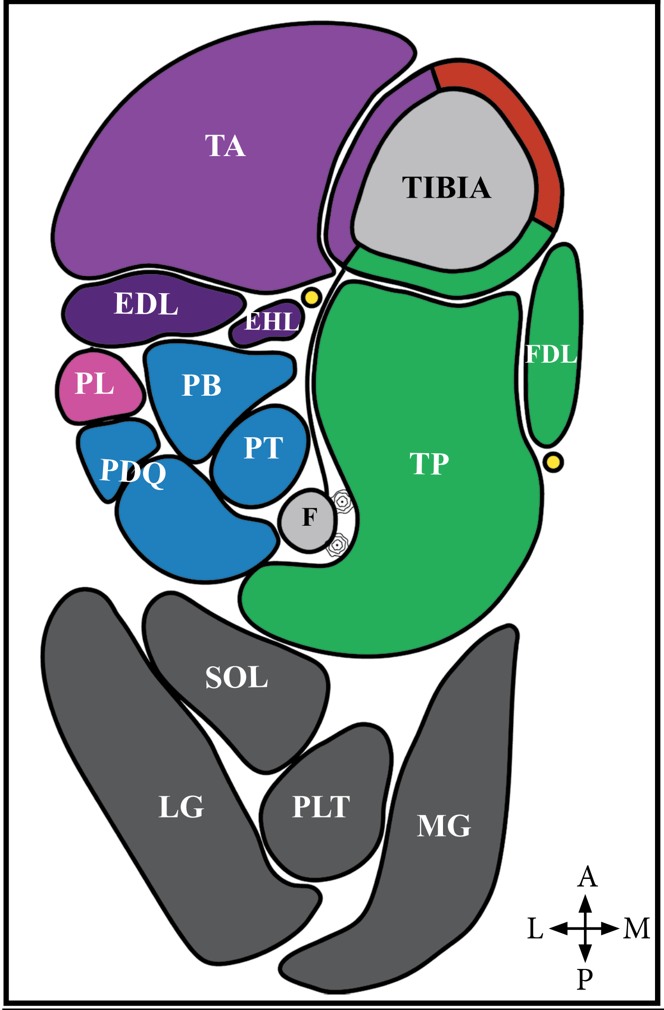


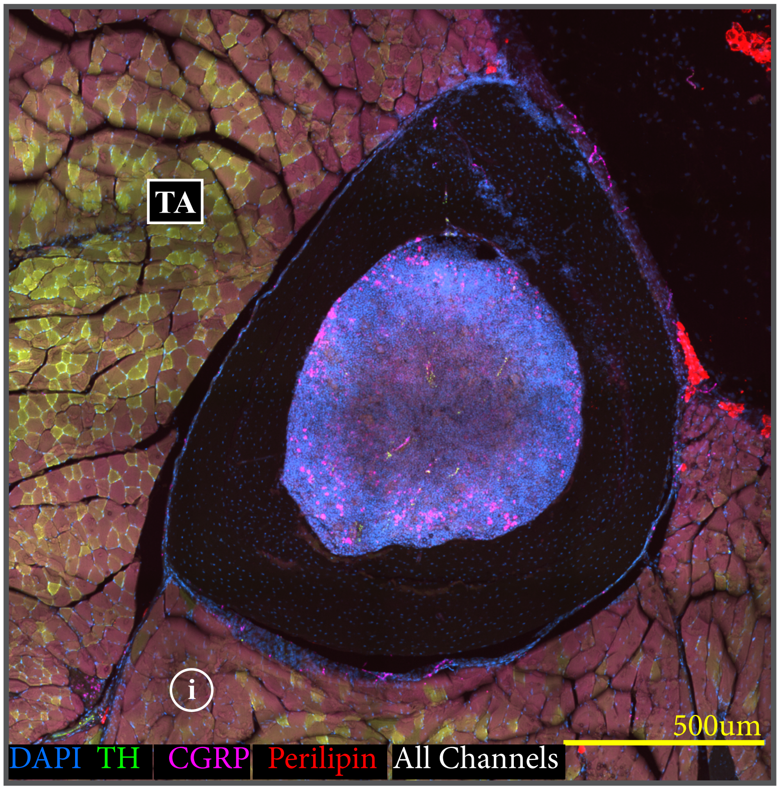


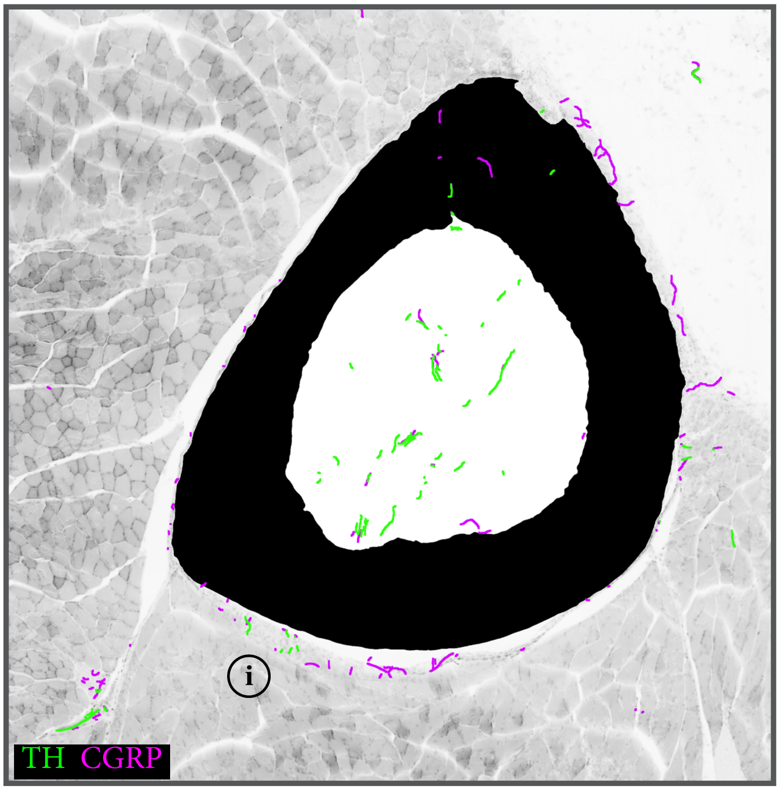


**Color Legend**

**Pattern Legend**

Hover over icon for more information

TEXT Hover over abbreviations for full definition

Use arrows to access keys and icons

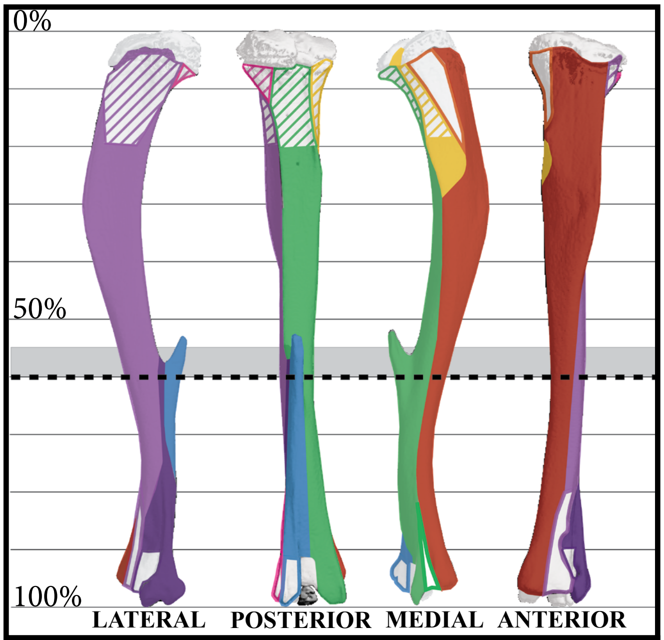


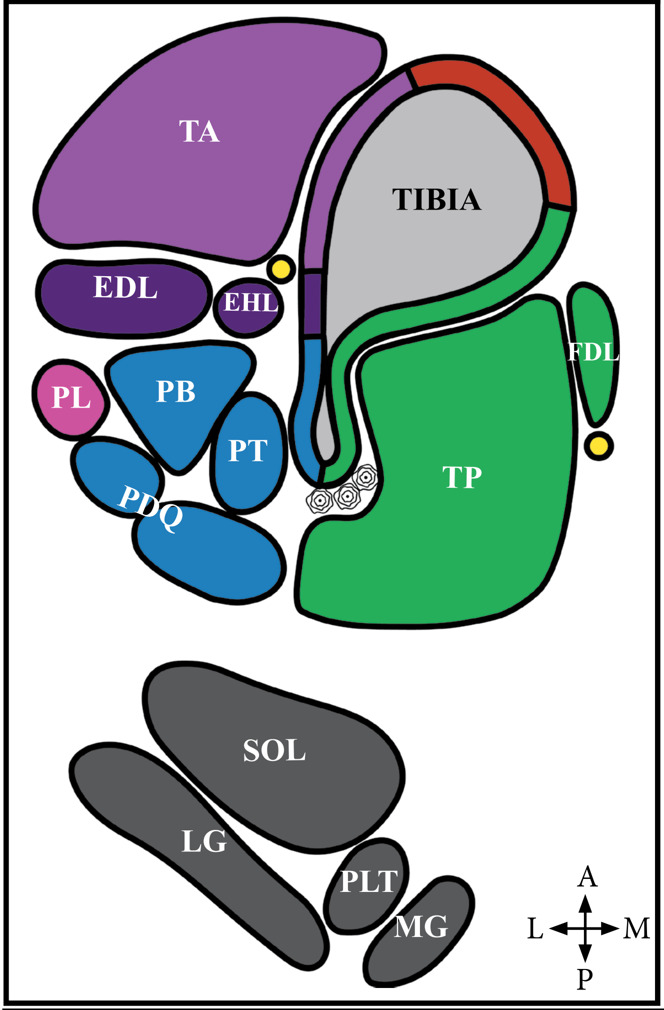


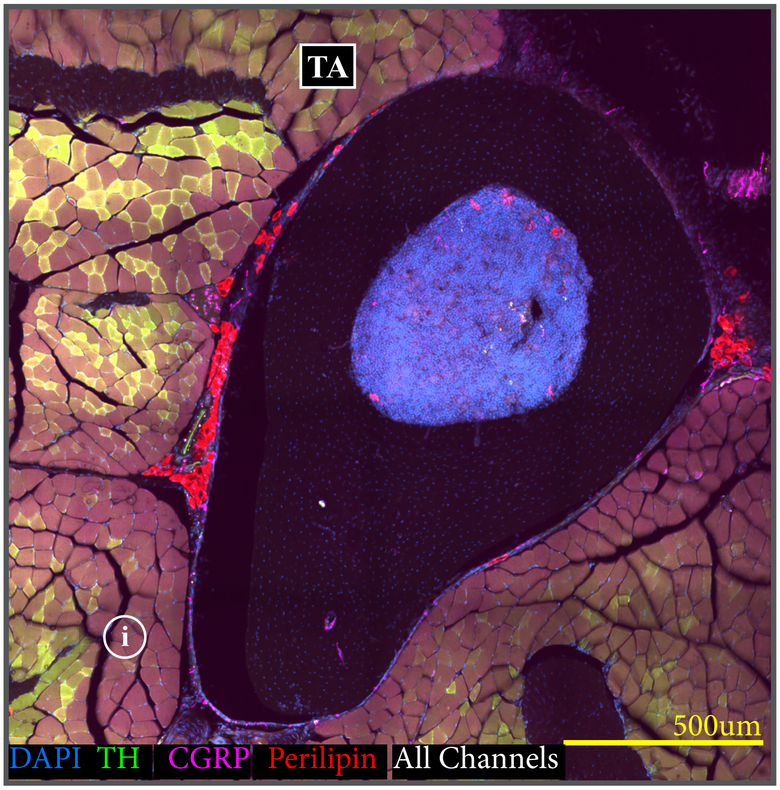


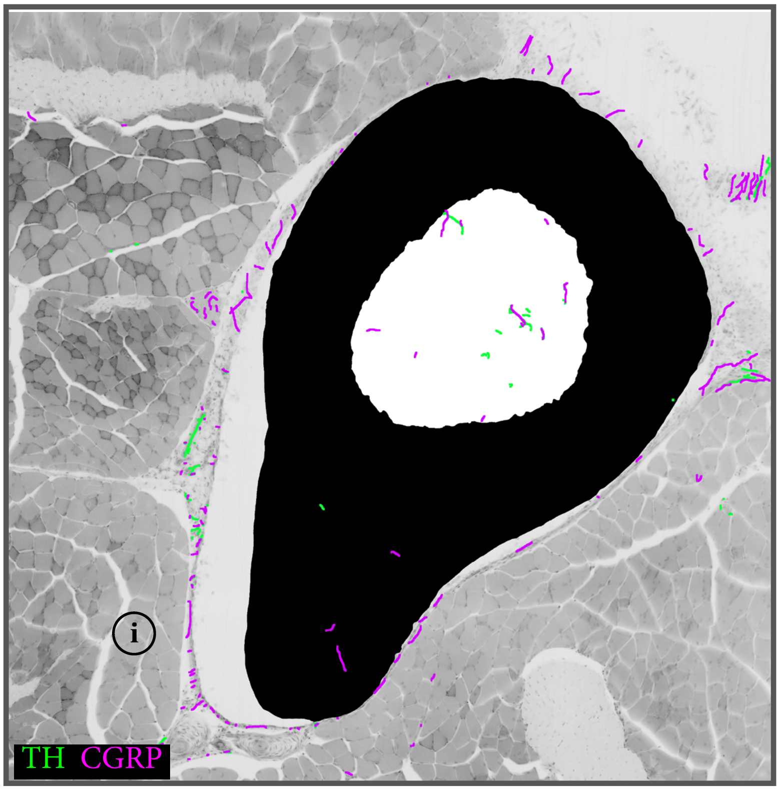


Color Legend

Pattern Legend

Hover over icon for more information

TEXT Hover over abbreviations for full definition

Use arrows to access keys and icons

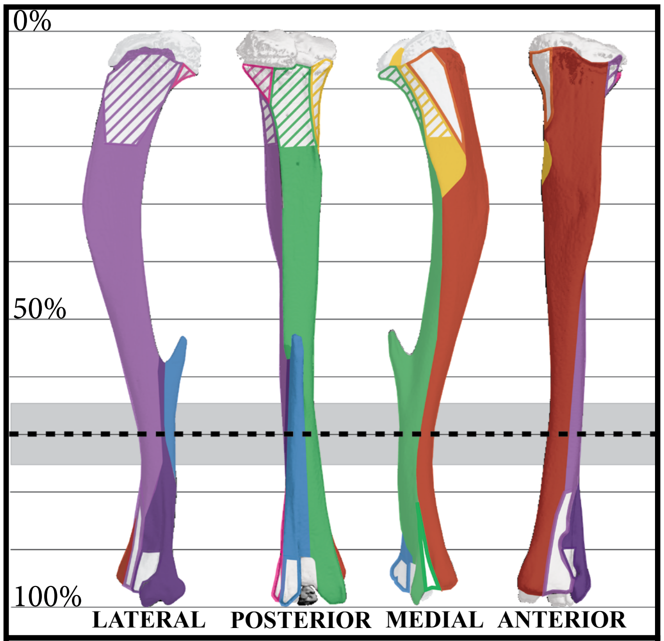


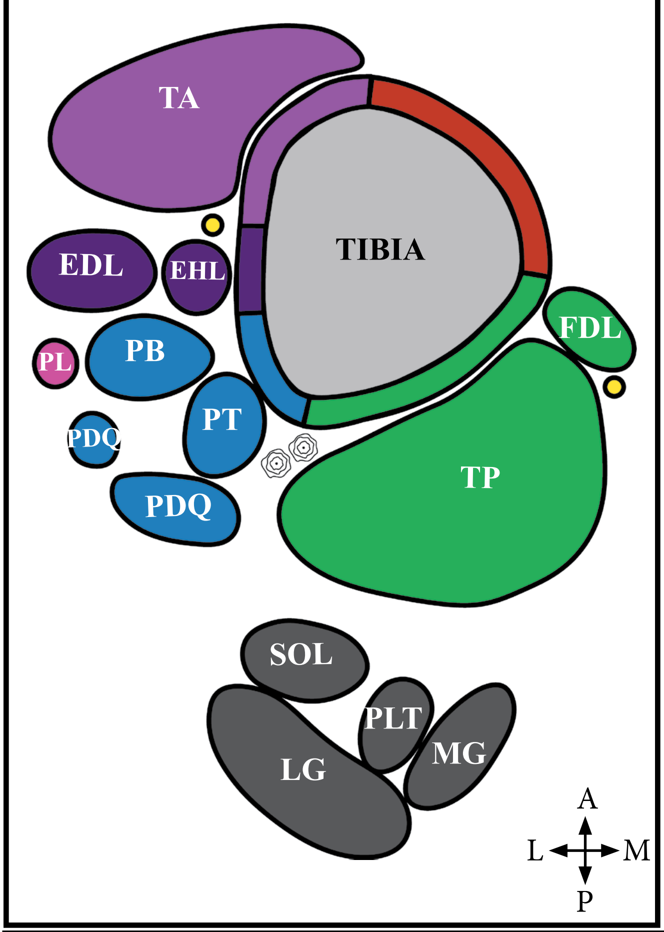


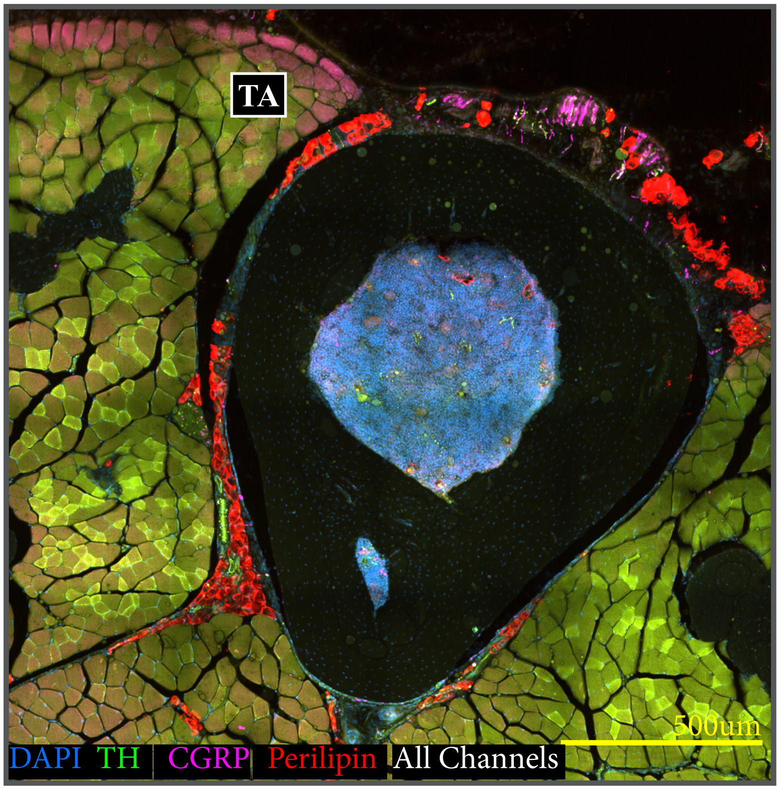


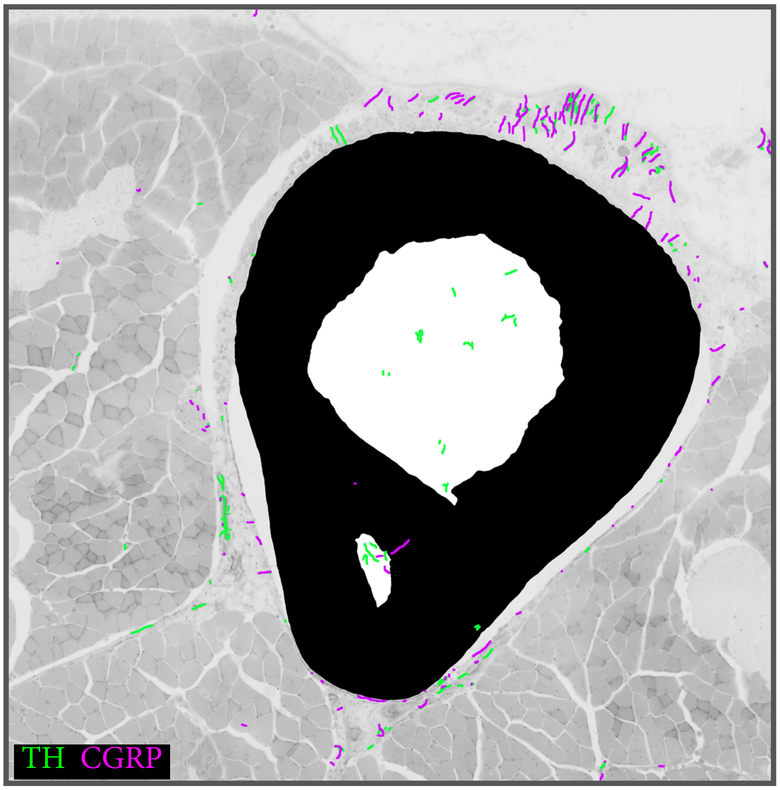


**Color Legend**

**Pattern Legend**

Hover over icon for more information

Hover over abbreviations for full definition

Use arrows to access keys and icons

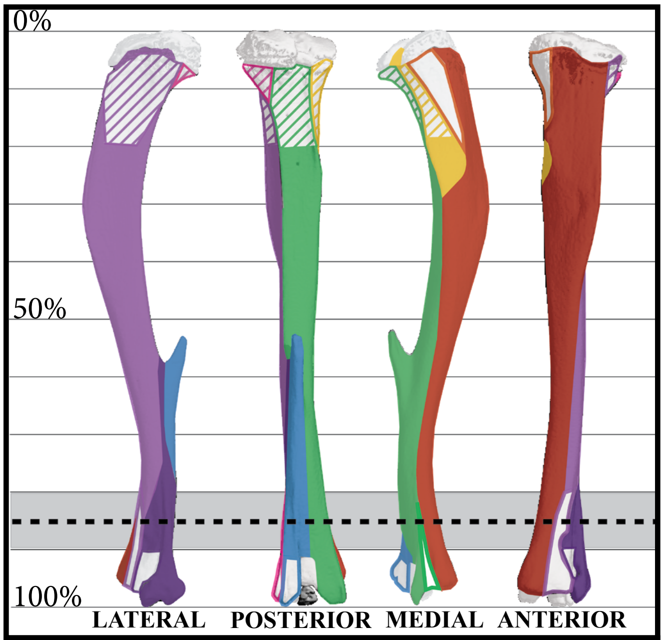


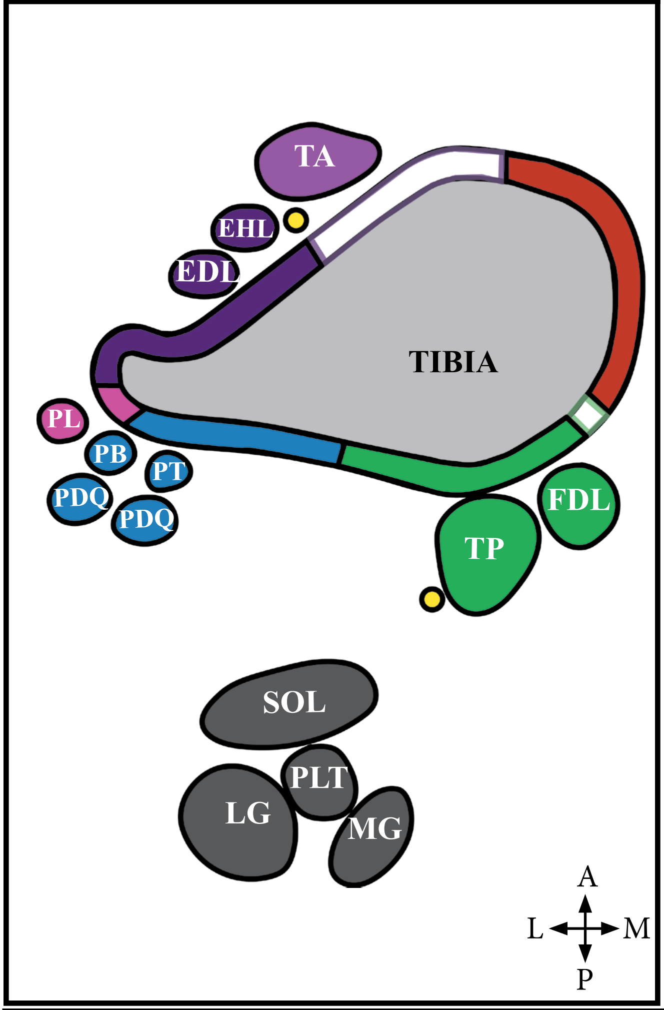


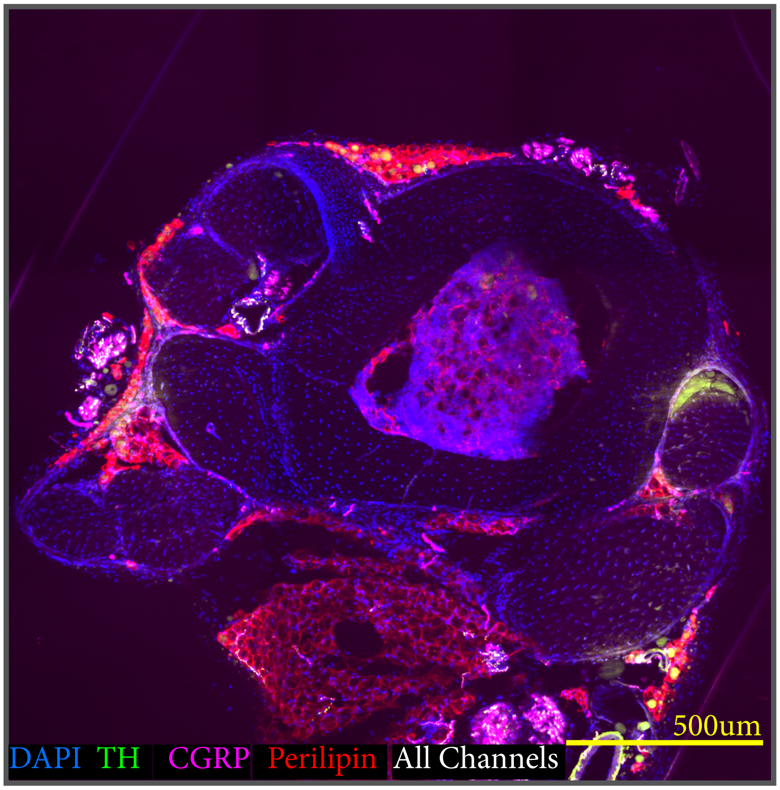


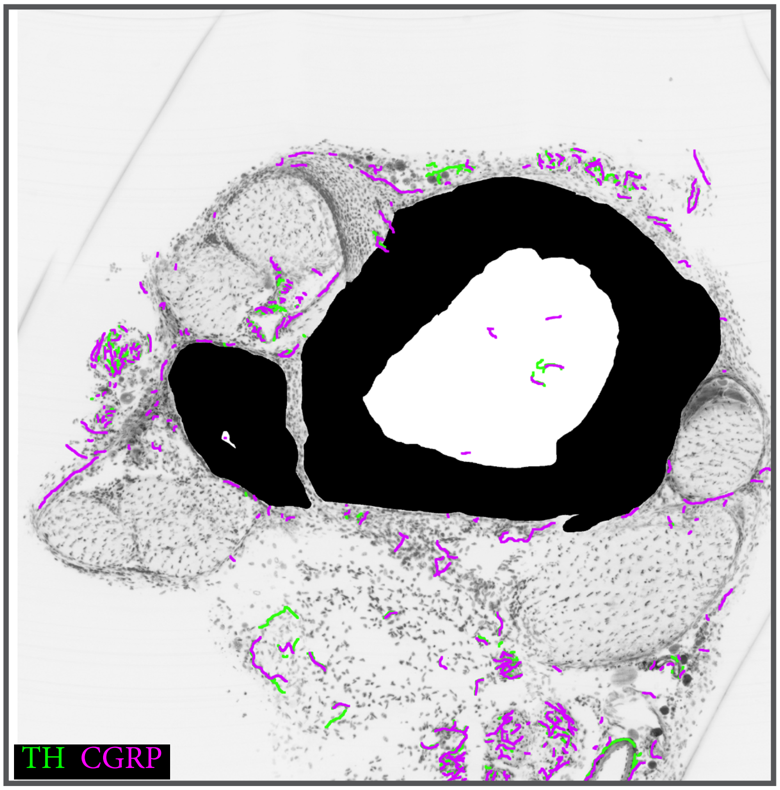


**Color Legend**

**Pattern Legend**

Hover over icon for more information

TEXT

Hover over abbreviations for full definition

Use arrows to access keys and icons

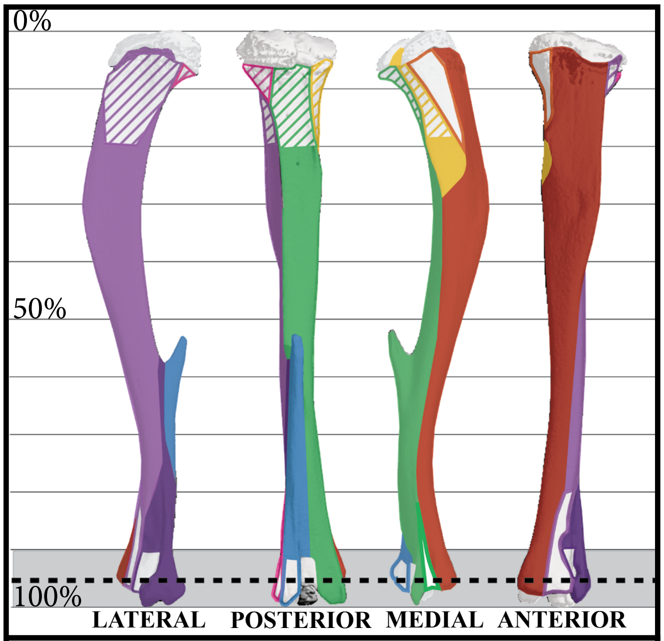


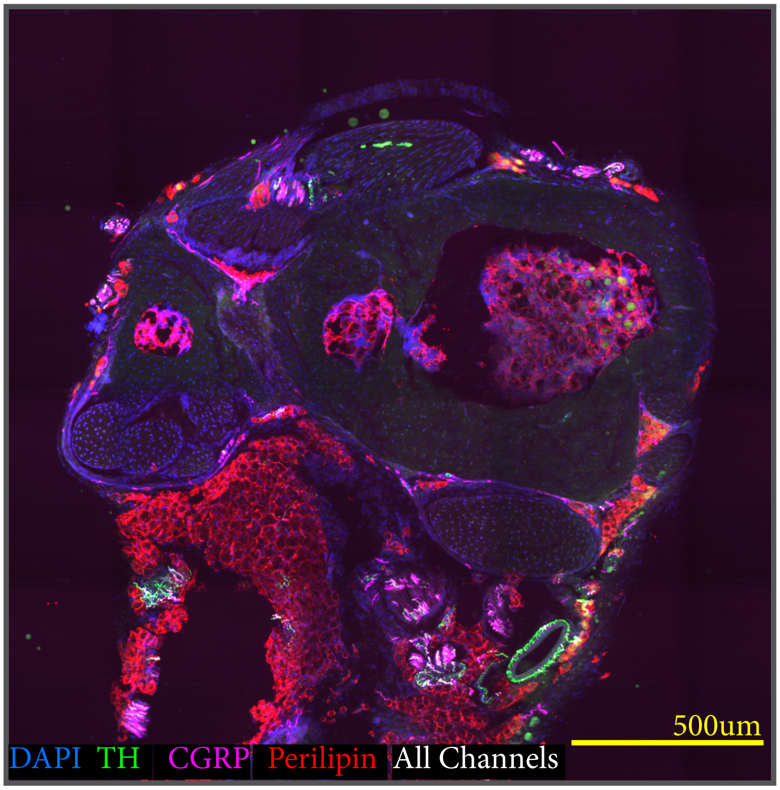


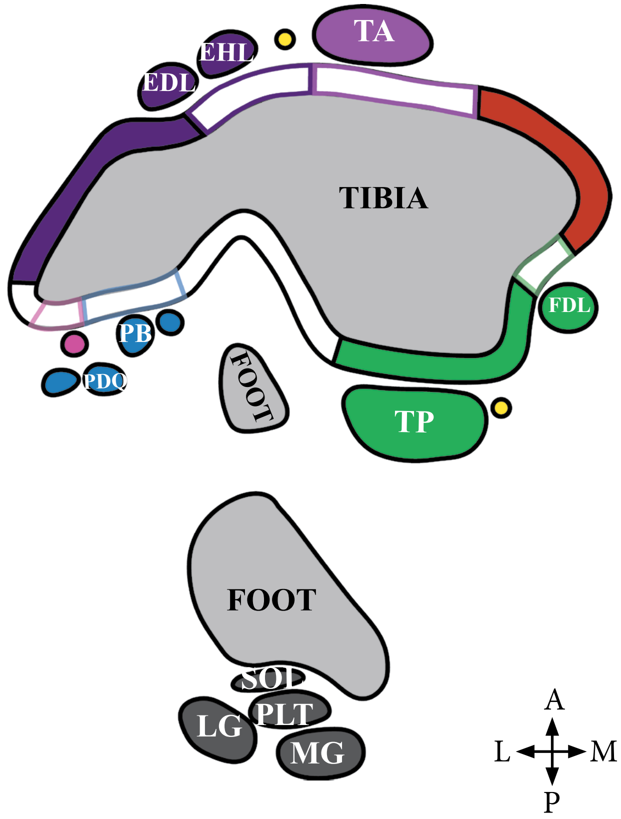


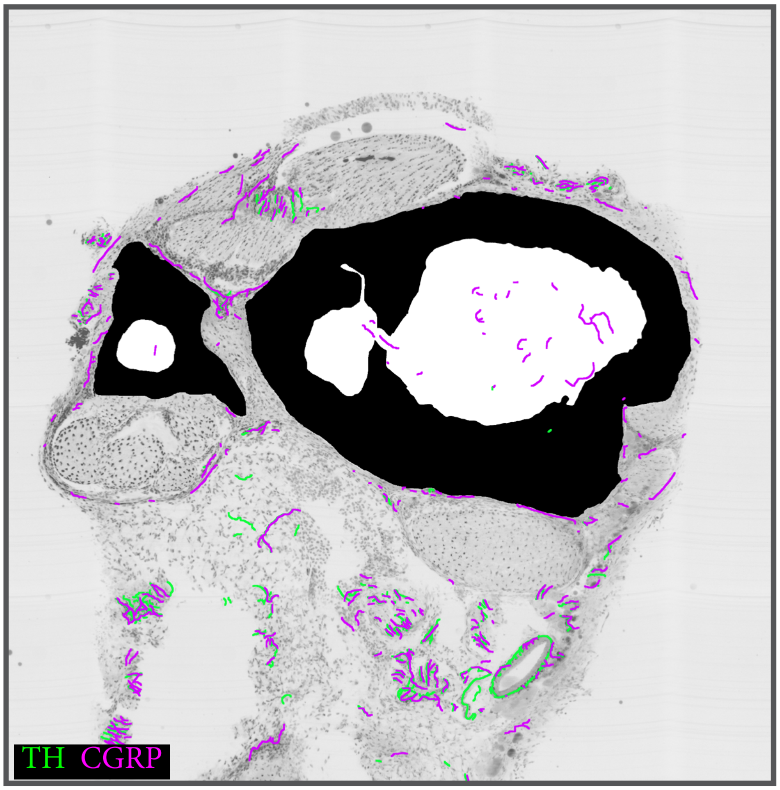


**Color Legend**

**Pattern Legend**

Hover over icon for more information

TEXT

Hover over abbreviations for full definition

Use arrows to access keys and icons
